# Chromosome-level genomics and historical museum collections reveal new insights into the population structure and chromosome evolution of waterbuck

**DOI:** 10.1101/2025.03.19.644014

**Authors:** Corey Kirkland, Xi Wang, Carla Canedo-Ribeiro, Lucía Álvarez-González, David Weisz, Alexandria Mena, Judy St Leger, Olga Dudchenko, Erez Lieberman Aiden, Aurora Ruiz-Herrera, Rasmus Heller, Tony King, Marta Farré

**Affiliations:** School of Natural Sciences, University of Kent, Canterbury, UK; Department of Biology, University of Copenhagen, Copenhagen, Denmark; Departament de Biologia Cellular, Fisiologia i Immunologia, Universitat Autònoma de Barcelona, Cerdanyola del Vallès, Spain; Institut de Biotecnologia i Biomedicina, Universitat Autònoma de Barcelona, Cerdanyola del Vallès, Spain; The Center for Genome Architecture, Baylor College of Medicine, Houston, USA; SeaWorld San Diego, San Diego, CA, USA; Cornell University College of Veterinary Medicine, Ithaca, NY, USA; The Center for Theoretical Biological Physics, Rice University, Houston, USA; The Aspinall Foundation, Port Lympne Reserve, Kent, UK; Durrell Institute of Conservation and Ecology (DICE), University of Kent, Canterbury, UK

**Keywords:** genome assembly, population genomics, historical DNA, chromosome evolution, Robertsonian fusions

## Abstract

Advances in the sequencing and assembly of chromosome-level genome assemblies has enabled the study of non-model animals, providing further insights into the evolution of genomes and chromosomes. Here, we present the waterbuck (*Kobus ellipsiprymnus*) as an emerging model antelope for studying population dynamics and chromosome evolution. Antelope evolutionary history has been shaped by Robertsonian (Rb) fusions, with waterbuck also showing variation in karyotype due to two polymorphic Rb fusions. These polymorphisms are variable between and within the two recognised subspecies, the common and defassa waterbuck. To provide new insights into waterbuck evolution, we firstly assembled a chromosome-level genome assembly for the defassa subspecies using PacBio HiFi and Hi-C sequencing. We then utilised museum collections to carry out whole genome sequencing (WGS) of 24 historical waterbuck skins from both subspecies. Combined with a previous WGS dataset (n = 119), this represents the largest study of waterbuck populations to date. We found novel population structure and gene flow between waterbuck populations and regions across the genome with high genomic differentiation between the two subspecies. Several of these regions were found around the centromeres of fixed and polymorphic Rb fusions, exhibiting signatures of low recombination and local population structure. Interestingly, these regions contain genes involved in development, fertility, and recombination. Our results highlight the importance of assembling genomes to the chromosome-level, the utility and value of historical collections in sampling a wide-ranging species to uncover fine-scale population structure, and the potential impacts of Rb fusions on genomic differentiation and the recombination landscape.

## Introduction

The waterbuck (*Kobus ellipsiprymnus*) is a large antelope species composed of two subspecies, the common (*K. e. ellipsiprymnus*) and defassa (*K. e. defassa*) waterbuck, with their taxonomic classification based on rump fur colouration, molecular data, and cytogenetics (Kingswood et al. 1998; Lorenzen et al. 2006; Wang et al. 2024). The species is distributed across central, eastern, and southern Africa, with the common subspecies found east of the Great Rift Valley, from Somalia in the north to South Africa in the south, while the defassa subspecies is found to the west of the Great Rift Valley, covering a larger distribution from Guinea Bissau in West Africa to Kenya and Tanzania in East Africa, as well as populations in Angola and Zambia in the south. The species is listed on the IUCN Red List as Least Concern and the defassa subspecies further listed as Near Threatened, due to population declines (Anon 2016), and therefore the species is becoming a conservation concern.

The two subspecies have high levels of genetic and genomic differentiation caused by a separation across the Great Rift Valley (Lorenzen et al. 2006; Wang et al. 2024). Although this separation occurred during the mid-Pleistocene, historical and recent gene flow has subsequently occurred across this geographic and climatic barrier (Wang et al. 2024). A hybrid zone exists in northern Tanzania and Kenya, where this admixture is suggested to be recent, with hybrid individuals reported with intermediate phenotypes (Lorenzen et al. 2006; Wang et al. 2024). Within each subspecies, population structure largely reflects geographic locality, with each subspecies further grouped into northern and southern population groups (Wang et al. 2024).

At the chromosome-level, waterbuck show inter- and intra-subspecies karyotypic variation in natural and captive populations due to the Robertsonian (Rb) fusion of two pairs of acrocentric chromosomes, syntenic to cattle (*Bos taurus*) chromosomes BTA6;18 and BTA7;11 (Kingswood et al. 1998; Kingswood et al. 2000; **Fig. 1A**). In the common waterbuck the BTA6;18 fusion is fixed, whilst the BTA7;11 fusion is polymorphic, resulting in three karyotypes: 2*n* = 52 (without the fusion), 2*n* = 51 (heterozygous for the fusion), or 2*n* = 50 (homozygous for the fusion). Whereas in the defassa, the BTA7;11 fusion is absent, and the BTA6;18 fusion has been found to be heterozygous (2*n* = 53) or absent (2*n* = 54). The remaining waterbuck karyotype is composed of three submetacentric autosomes, formed from historical Rb fusions that have become fixed within the species (syntenic to cattle chromosomes BTA1;19, BTA2;25, and BTA5;17), a metacentric X chromosome, an acrocentric or submetacentric Y chromosome, and 19-23 acrocentric autosomes (Kingswood et al. 1998; Kingswood et al. 2000).

**Figure 1:**
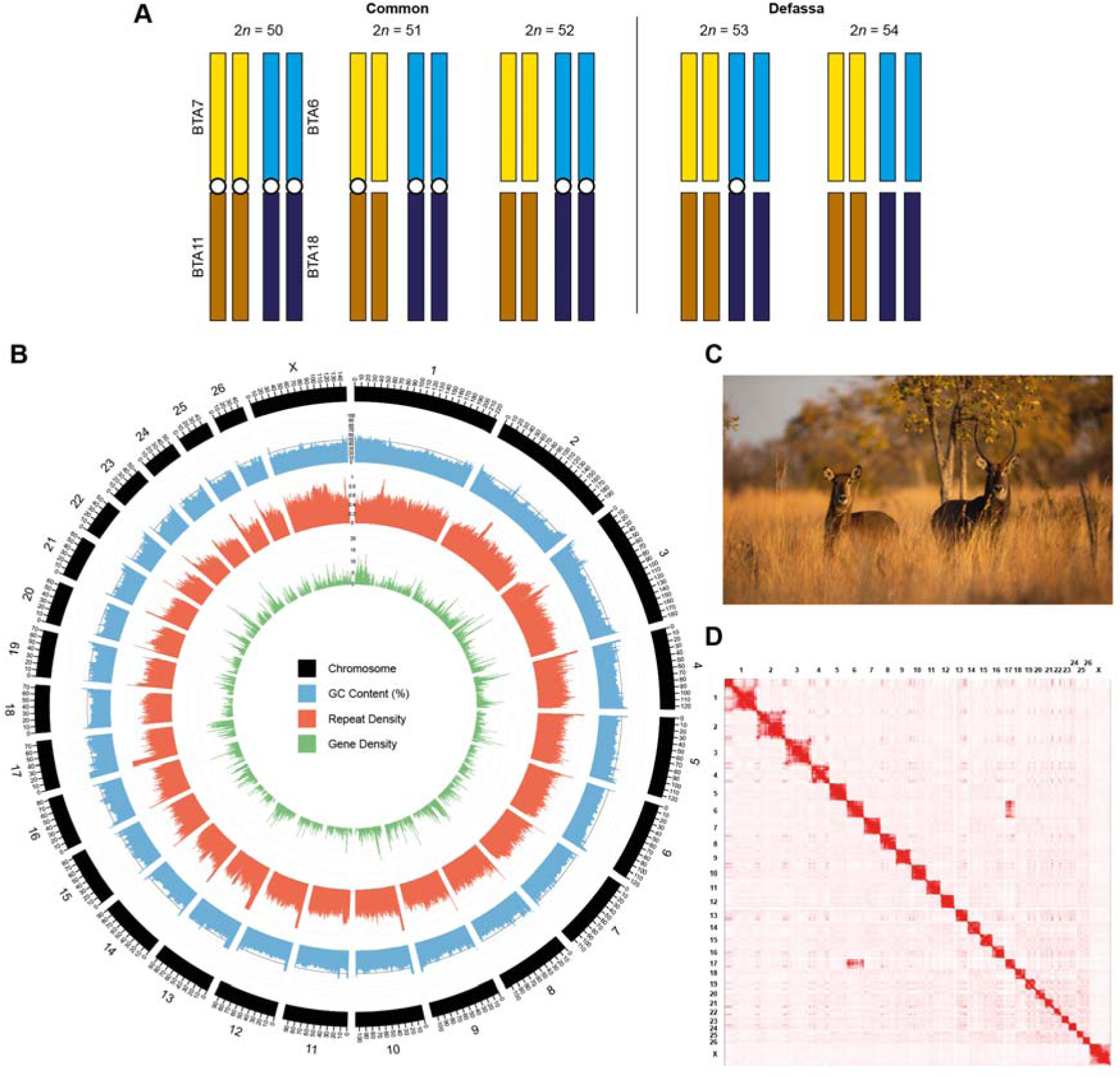
Chromosome-level genome assembly generated for the defassa waterbuck (2*n* = 54). (A) Ideogram of the previously reported karyotypes in waterbuck (Kingswood et al. 1998) with the two polymorphic Robertsonian fusions shown (homologous to cattle BTA6;18 and BTA7;11). Chromosomes are coloured by homology to cattle chromosomes and Rb fusions are depicted by a white circle. (B) Circos plot of waterbuck chromosomes, with GC content (%), repeat density, and gene density calculated in 100 Kb windows. (C) A female and male waterbuck. (D) Hi-C interaction matrix for the 2*n* = 52 sample mapped to the chromosome-level genome assembly (2*n* = 54), showing interchromosomal interactions between KEL6 and KEL17 (syntenic to cattle BTA6 and BTA18, respectively), as a result of the Rb fusion.

Waterbuck population dynamics have recently been studied at the genomic-level (X. Wang et al., 2024) using a scaffold-level short-read assembly for the waterbuck as reference (Chen et al. 2019), as well as the chromosome-level genome of goat (*Capra hircus*). However, the previous waterbuck reference genome is composed of more than 88,000 scaffolds, with an N50 of 0.78 Mb (Chen et al. 2019). As ruminants have highly repetitive genomes and several well characterised chromosome rearrangements (Arias-Sardá et al. 2023), and waterbuck show karyotypic variation, a chromosome-level genome is required for this species to uncover the consequences of these chromosome rearrangements and to further understand the evolution of the species. Moreover, the previous population genomic study (Wang et al. 2024) particularly focused on populations near the contact zone between the two subspecies, but a wider genomic sampling is required for this species to uncover fine-scale population structure, due to its large distribution across Africa.

Here, we sequenced and assembled a chromosome-level genome for the waterbuck, achieving high contiguity and completeness. We then generated genomic sequences for 24 historical museum samples and integrated this with 119 previously published modern samples (Wang et al. 2024). This allowed us to study population structure at both a broader and a more finely resolved scale than has previously been reported. Using the newly generated high-quality genome assembly, we investigated genomic differences at the chromosome-level between the two subspecies of waterbuck. Our analysis revealed putative signatures of Rb fusions in waterbuck, and other novel variation across the genome. This study provides a greater understanding of population structure and chromosome rearrangements in shaping subspecies divergence in waterbuck with potential implications for the conservation management of this antelope.

## Results and Discussion

### Sequencing and assembling a chromosome-level genome for the waterbuck

In order to explore population structure and chromosome evolution within the waterbuck at the chromosome level, a high-quality genome was required. To do this, we firstly established a cell culture of a female defassa waterbuck. Karyotyping revealed that the sample had a standard karyotype for the defassa of 2*n* = 54, without any of the two polymorphic Rb fusions (**Supplementary Fig. S1**). The sample was then sequenced with PacBio HiFi (21X coverage, read N50: 18.89 Kb) and assembled to contig-level (3.15 Gb, 1,071 contigs, N50: 71.17 Mb, L50: 17; **Supplementary Fig. S2A**). The N50 of the contig-level genome assembly was 343 times greater than the previous short-read assembly of 0.21 Mb (Chen et al. 2019), demonstrating the power of highly accurate PacBio HiFi long-reads in increasing contiguity.

To scaffold the genome to chromosome-level, Hi-C data was generated and sequenced from the defassa sample (2*n* = 54) and an additional female common waterbuck sample (2*n* = 52; **Supplementary Fig. S3A and S3B**). Due to better data quality the Hi-C scaffolding was completed using data from the common waterbuck sample (2*n* = 52). Hi-C data from the defassa waterbuck sample was used to help resolve the genome assembly karyotype to match the defassa subspecies (2*n* = 54), and this was confirmed with synteny to cattle chromosomes (**Supplementary Fig. S3A and S3D**) and the previous cytogenetic study (Kingswood et al. 1998). In this study, we also further curated the genome by reorienting and reordering chromosomes by size (**Supplementary Table S1**). This resulted in a final curated genome assembly for the defassa waterbuck (**Fig. 1B and 1D, Supplementary Fig. S2B and S3**). The chromosome-level genome was highly contiguous (3.15 Gb, 1,014 scaffolds, N50: 98.45 Mb, and L50: 12) containing 26 autosomes and one sex chromosome (**Fig. 1D**). This improved the N50 of the previous scaffold-level assembly (0.78 Mb; Chen et al., 2019) by 125 times. The final assembly presents 96.27% of BUSCO mammalian genes, with 94.41% being single copy (**Supplementary Fig. S2B**). This meets the VGP-2020 standards (a contig NG50 greater than 10 Mb, a scaffold NG50 equal to the chromosome NG50, and gene completeness greater than 95%; Rhie et al. 2021). Our genome assembly is therefore highly contiguous and complete, and compares to other ruminant genomes assembled with the same methodology, such as the takin (*Budorcas taxicolor*) genome with a contig N50 of 68.05 Mb, scaffold N50 of 101.27 Mb, and a BUSCO completeness of 94.2% (Li et al. 2023). The mitochondrial genome was also assembled from the PacBio HiFi reads, generating a genome of 16,427 bp in length including 37 genes, of which 13 were protein coding, 22 were tRNA, and two were rRNA (**Supplementary Fig. S4)**, in line with other sequenced bovids (Yang et al. 2013).

A total of 24,645 protein-coding genes were predicted using homology-based annotation with cattle (*Bos taurus*; ARS-UCD2.0) and goat (*Capra hircus*; ARS1.2) as references (**Fig. 1B and Supplementary Table S2**), similar to other ruminant genomes (21,667 in cattle and 20,663 in goat). Around 54.80% (1.73 Gb) of the genome was repetitive, with LINEs representing the majority of repeats (24.26%), followed by satellite or centromeric repeats (14.24%), and SINEs (9.70%; **Supplementary Table S3**). RTE-BovB elements made up the majority of LINEs (11.02%) and are characteristic of bovid genomes, where they have expanded since they first occurred in the ancestor of ruminants through horizontal gene transfer (Adelson et al. 2009; Ivancevic et al. 2018). GC content (%) was highest at the start and ends of chromosomes, while repeat density was highest in regions surrounding the expected locations of centromeres (**Fig. 1B**).

### Population genomics in waterbuck using modern and historical samples

Previous studies on the population genetics and genomics of the waterbuck have found that genetic structure reflects the geographic localities of populations, and that gene flow between the two subspecies is both historical and ongoing (Lorenzen et al. 2006; Wang et al. 2024). To further understand the evolution of waterbuck and provide a wider sampling distribution across the species range we utilised museum collections. In total we sampled 24 historical waterbuck skins (1900-1938), extracted DNA, and performed WGS at low coverage (∼5X; **Fig. 2A and Supplementary Table S4**). Of the historical samples, 20 were taxonomically classified as the defassa subspecies (from locations in Angola, Cameroon, Chad, Democratic Republic of Congo; DRC, Guinea Bissau, South Sudan, and Tanzania) and four as common (from locations in Ethiopia and Somalia). We also combined our historical data with 119 modern WGS data from the previous genomic study (X. Wang et al., 2024; **Fig. 2A and Supplementary Table S4**). Modern data included six populations classified as defassa from Uganda (QENP and KVNP), Zambia (Kafue), Tanzania (Maswa and Ugalla), and Ghana (Samole), and four common waterbuck populations from Zimbabwe (Matetsi), Zambia (Luangwa), and Kenya (Samburu and Nairobi).

**Figure 2:**
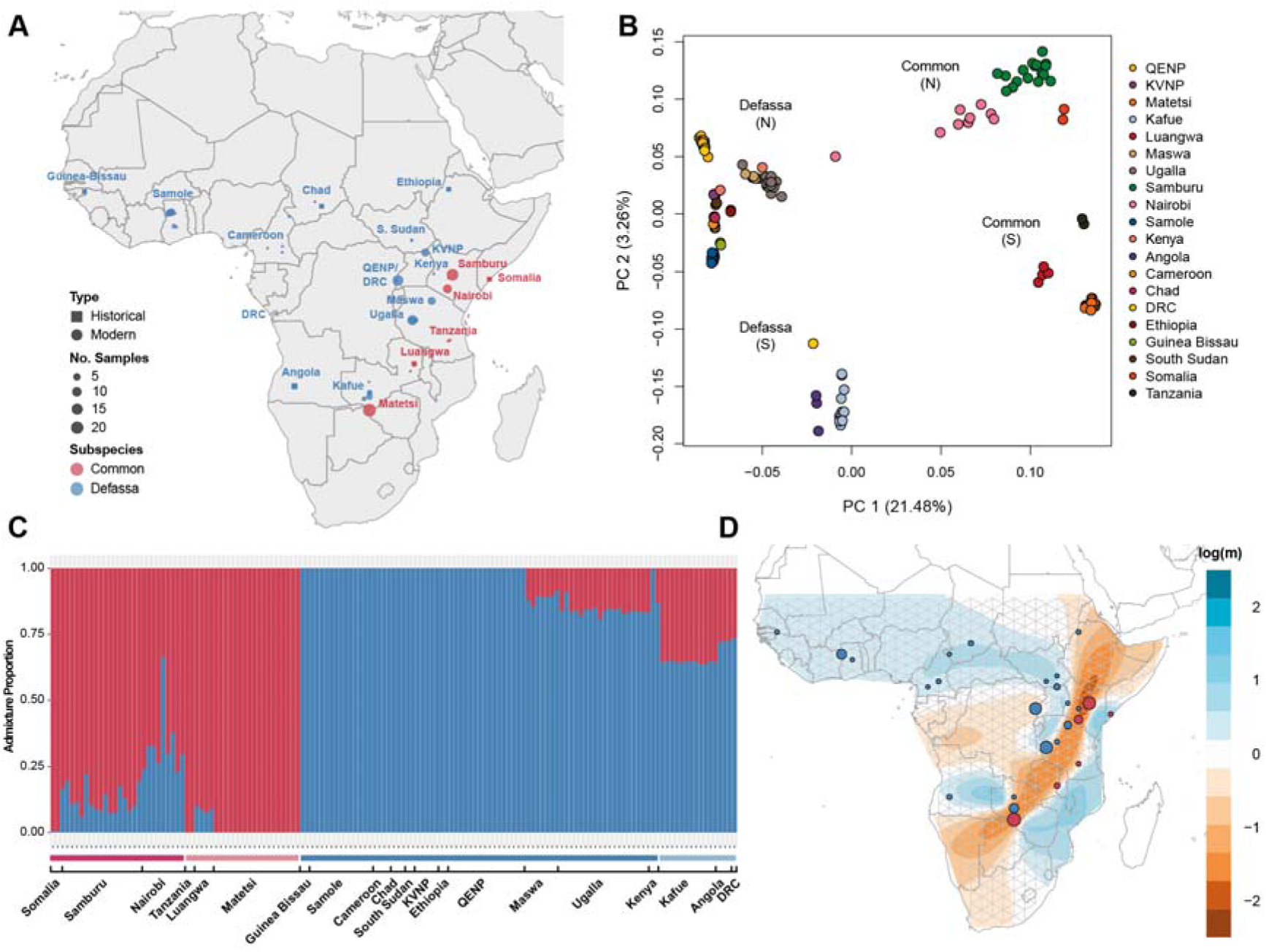
Population structure in the waterbuck. (A) Sampling map of the historical and modern samples. (B) PCA using all genomic sites grouped by population showing the North (N) and South (S) population groups for each subspecies. (C) Admixture proportions at K=2 grouped by populations, subspecies, and geographical location. Red indicates common waterbuck and blue defassa waterbuck, with a darker shade representing the North group and a lighter shade the South group. (D) EEMS log migration rates (*m*) between populations, with common populations indicated by red circles and defassa blue circles. The size of the circles represents the number of samples at a given location.

All WGS data was mapped to the waterbuck chromosome-level genome assembly, with an average coverage of 4.03X across the 143 samples (**Supplementary Table S4**). To produce a set of high confidence SNPs and genomic areas for downstream analyses, we conservatively filtered genomic sites, by removing repetitive regions, positions with excess heterozygosity, low mappability, and regions with low or high depth of coverage. This resulted in 40.19 % of the reference genome available for our analyses (**Supplementary Table S5**).

As historical samples are prone to DNA damage over time, we then estimated DNA damage of the mapped sequencing reads (**Supplementary Fig. S5**). C to T substitutions were highest at the starts of paired end reads, with sample ‘WB_1b_5X’ having the highest rate (0.026) and sample “WB_2h_5X” the lowest (0.006). Whereas G to A substitutions were highest at the end of collapsed reads (0.090 to 0.044). To understand the impact of this potential DNA damage, we calculated genome-wide heterozygosity on all genomic sites and compared this to transversion sites (**Supplementary Fig. S6**). The historical samples had a higher heterozygosity than modern samples when using all sites (p = 0.0001), but no significant difference when using only transversion sites (p = 0.8271), suggesting some differences due to historical DNA damage, rather than a change in heterozygosity due to any population decline. The DRC sample ‘WB_3k_5X’ had extremely high levels of heterozygosity (0.0129 for all sites and 0.0054 for transversion sites) and therefore was removed from the EEMS and F_ST_ analyses. Focusing on transversion sites for historical and modern populations, Luangwa had the highest average heterozygosity of all populations (0.0017), whereas Angola, Cameroon, and KVNP had the lowest average heterozygosity (0.0009; **Supplementary Fig. S6**).

Principal Component Analysis (PCA), admixture proportions (at K = 2, K = 3, K = 4, and K = 12 estimated populations), and overall F_ST_ were computed on genomic sites across the 26 autosomal chromosomes and 143 samples. Overall, PC1 (21.48%) of the PCA showed a split between the two subspecies, with the exception of one sample from Nairobi, which had previously been shown to be recently admixed (Wang et al. 2024), while PC2 (3.26%) grouped populations in the north and south separately (**Fig. 2B**). A similar PCA result was found using only transversion sites, confirming that historical samples were not causing differences in genomic variability due to DNA damage at transition sites (**Supplementary Fig. S7**). Moreover, the historical samples grouped with the modern samples by geographic locality, suggesting that population structure has been maintained over the past 100 years.

Population structure was also supported by moderate genomic differentiation (F_ST_ = 0.214) between the two subspecies; and between northern and southern groups in the common and defassa populations (F_ST_ = 0.115 and 0.139, respectively; **Supplementary Fig. S8**). Admixture at K = 2 grouped individuals into the two subspecies, with varying proportions of admixture (**Fig. 2C**), at K = 3 defassa were split into two populations and common into one, at K = 4 defassa were split into three populations and common into one, and at K = 12 individuals were grouped more similarly to their geographic locality and populations (**Supplementary Fig. S9**).

Taking a closer look at each population group, the Defassa North populations in West and Central Africa (Guinea Bissau, Samole, Cameroon, Chad, and South Sudan) and East Africa (Ethiopia, KVNP, and QENP) grouped together on the PCA and admixture analyses, showing no signs of admixture with common populations (**Fig. 2 and Supplementary Fig. S9**). Whereas populations of Defassa North closer to the Great Rift Valley were clustered together on the PCA, but showed some admixture with common (Maswa, Ugalla, and the historical Kenyan sample closer to the common waterbuck population of Samburu), confirming that admixture was isolated to the contact zone between the two subspecies in this region.

The Defassa South group (Kafue, Angola, and the southern DRC individual) clustered between the defassa and common groupings on PC1 of the PCA, and separately on PC2, with the DRC individual clustered further towards the defassa populations of Samole and Guinea Bissau in the northwest on PC1 (**Fig. 2B**). These populations also had varying degrees of admixture proportions with common at K = 2 (**Fig. 2C**). This was surprising for populations in Angola and DRC which are not geographically close to the contact zone of the two subspecies. At K = 3, the Defassa South population had admixture with defassa in central Africa and common waterbuck, and at K = 4 grouped separately (**Supplementary Fig. S9**). There was also some admixture with the Defassa South group in Maswa at K = 4. These results could suggest there has historically been movement of defassa waterbuck both around the east and west of the Congolese Rainforest.

Populations of the Common North group showed varying proportions of admixture with defassa, with animals from Nairobi displaying the largest admixture proportions with an average of 0.34 across all nine animals at K = 2 (**Fig. 2C**), supporting a well-known hybrid zone in this region in Kenya (Lorenzen et al. 2006; Wang et al. 2024). Interestingly, the Common South population of Matetsi is in close proximity to Kafue (Defassa South) but showed no admixture with defassa at K = 2 (**Fig. 2C**), whereas the population of Luangwa displayed admixture proportions between 0.07 and 0.10. This suggests that gene flow is occurring between Luangwa and the Defassa South group, but not between Matetsi and the Defassa South group. Interestingly, this is partially supported by a previous TreeMix analysis, where a migration event from Luangwa to Kafue was found (Wang et al. 2024); however, our results might indicate a bidirectional migration event in this region. Our results point to a barrier to gene flow between Matetsi and the Defassa South group, with the Zambezi River potentially acting as a geographic barrier between the two populations. No admixture was found in the far east in the historical population of Tanzania. Our results support the previous genomic study in finding regions along the contact zone with recent admixture (Wang et al. 2024), but we also uncover novel population structure in waterbuck distributed more geographically distant from the contact zone, such as in the southern defassa populations of Angola and DRC.

Estimated Effective Migration Surface (EEMS) was used to visualise gene flow and genetic diversity between populations (**Fig. 2D and Supplementary Fig. S10**). A strong barrier to gene flow was found across the subspecies divide, reflecting the Great Rift Valley (**Fig. 2D and Supplementary Fig. S10**). Decreased gene flow was also found around the Congolian Rainforest, creating an historical barrier between northern and southern defassa populations, which extends towards the Great Rift Valley (**Fig. 2D**). Mean diversity rates were highest between populations in East Africa around the Great Rift Valley (Ugalla, Maswa, Nairobi, Samburu, Somalia, and Luangwa), whilst lowest in the northwest (Guinea Bissau and Samole) and Ethiopia (**Supplementary Fig. S10**). The additional historical samples in central and southern Africa resulted in a better fit of reduced gene flow across the Great Rift Valley and the Congolian Rainforest than previously reported (Wang et al. 2024), and a more finely resolved and complete overview of the geographical barriers to gene flow in the waterbuck. These results support the present differentiation between the two subspecies and the north/south groupings in waterbuck. It also demonstrates the utility of museum collections in enabling the wider sampling of species with large distributions and detecting fine-scale population structure and gene flow.

### Genomic variation between waterbuck subspecies

As waterbuck showed substantial population structure between the two subspecies, we then investigated which regions of the genome were highly differentiated, using the newly assembled chromosome-level reference genome to place this into the context of waterbuck chromosomes (KEL). F_ST_ was estimated in 10 Kb windows containing greater than 1,000 sites after filtering (**Fig. 3**) and a total of 242 windows were in the 99.9^th^ percentile (F_ST_ ≥ 0.671). Overlapping these windows were 104 annotated protein-coding genes (**Supplementary Table S6**). The highest F_ST_ window containing annotated genes was on chromosome KEL21 (F_ST_ = 0.770) containing the gene MTMR14, encoding for a myotubularin related protein that plays a role in autophagy (Gibbs et al. 2010). This was followed by a window on KEL13 (F_ST_ = 0.760) containing the gene RTEL1, encoding a DNA helicase that maintains the integrity of telomeres and the stability of the genome by dissembling DNA secondary structures, facilitating DNA replication, repair, and recombination (Vannier et al. 2014). Knocked-out mice for this gene have been shown to have reduced cell proliferation, chromosomal fusions, and heterogeneity of telomere lengths (Ding et al. 2004).The absence of RTEL1 in the nematode *Caenorhabditis elegans*has been shown to cause complex chromosomal rearrangements as the gene prevents heterologous recombination during meiosis (León-Ortiz et al. 2018). Given the role of RTEL1 in maintaining telomere homeostasis and genome integrity, and the importance of telomeres in the formation of Rb fusions, it may therefore play a role in the polymorphic chromosome fusions in waterbuck.

**Figure 3:**
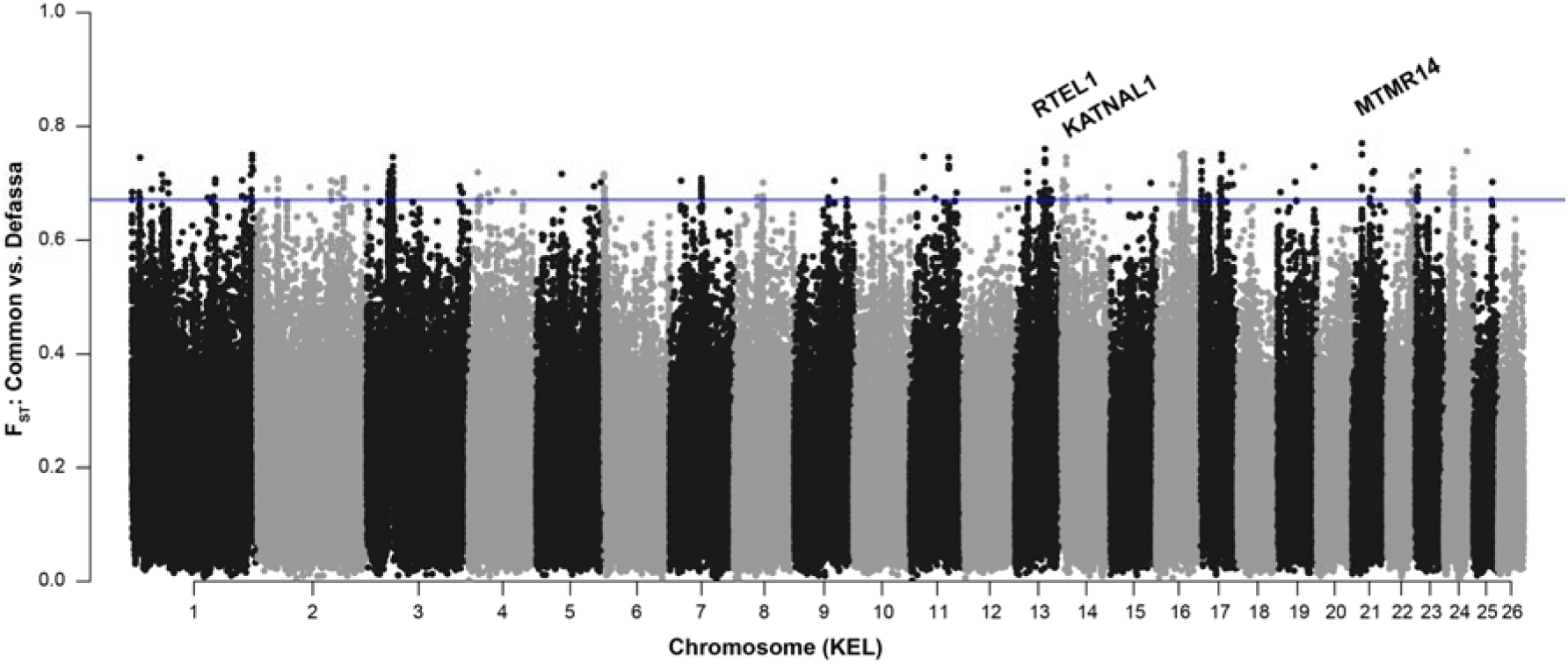
Genomic differentiation (F_ST_) calculated in 10 Kb windows between the common and defassa subspecies across waterbuck chromosomes (KEL). The blue line represents the 99^th^ percentile of F_ST_ windows.

Moreover, additional regions with high levels of genomic differentiation contained genes involved in chromatin (KMT2C, PPP1CA/PPP1CB, and BPTF), microtubules (STMN2, KATNAL1, KATNA1, and RABGAP1), and embryogenesis (LEUTX and HELZ). Relevant to reproduction, the gene KATNAL1 is involved in spermiogenesis, regulating chromosome alignment, segregation, and cytokinesis during male meiosis. Its loss of function causes the disruption of microtubules, leading to male infertility (Smith et al. 2012). When analysing the Gene Ontology (GO) terms of the 104 genes located in highly differentiated regions, we found these genes were statistically overrepresented for five Biological Process GO (negative regulation of t-circle formation, linoleic acid metabolic process, hepoxilin biosynthetic process, and lipoxygenase pathway) and three Molecular Function GO terms (RNA polymerase II binding, phosphatidylinositol activity, and linoleate activity; **Supplementary Table S7**).

### Putative signatures of chromosome rearrangements in waterbuck

Previous genetic and genomic studies in waterbuck (Lorenzen et al. 2006; Wang et al. 2024) have been unable to assess the impacts of Rb fusions on genomic differentiation. Using our new chromosome-level genome, we next explored the chromosomes involved in fixed (KEL1, KEL2, and KEL3) and polymorphic (KEL6;17 and KEL8;9) Rb fusions in more detail. We identified several regions of high F_ST_ between the two subspecies near the centromeres of chromosomes involved in fixed and polymorphic Robertsonian fusions within waterbuck.

Contrary to the two other submetacentric chromosomes (KEL1 and KEL2), we identified two blocks of high F_ST_ in the pericentromeric region of KEL3, a fixed Rb fusion in waterbuck (and fixed in the genus *Kobus*), homologous to cattle chromosomes BTA2 and BTA25 (**Fig. 4A and Supplementary Fig. S12**). These blocks were defined as large regions with elevated F_ST_ compared to adjacent genomic sites and were found on chromosome KEL3 between 38,425,000-41,405,000 bp and 45,415,000-47,265,000 bp (99.9^th^ percentile of the F_ST_ windows in the region was 0.719 and 0.744, respectively). These pericentromeric regions are in high linkage disequilibrium (LD) across the centromeric region, but only in the common waterbuck subspecies, suggesting that recombination is suppressed in this entire region (**Fig. 4C, 4D**, and **Supplementary Fig. S11**). Local PCAs of each of these two regions resulted in three groupings on PC1 (61.440% and 65.620% variance; **Fig. 4B and 4D**), with a different structure than the genome-wide PCA (**Fig. 2B**). The PCA on the KEL3 region grouped defassa populations on the left (and one common Nairobi sample), common on the right (and two defassa Ugalla samples), and individuals from populations of both subspecies near the contact zone (defassa: Ugalla, common: Nairobi, and Samburu) in the centre.

**Figure 4:**
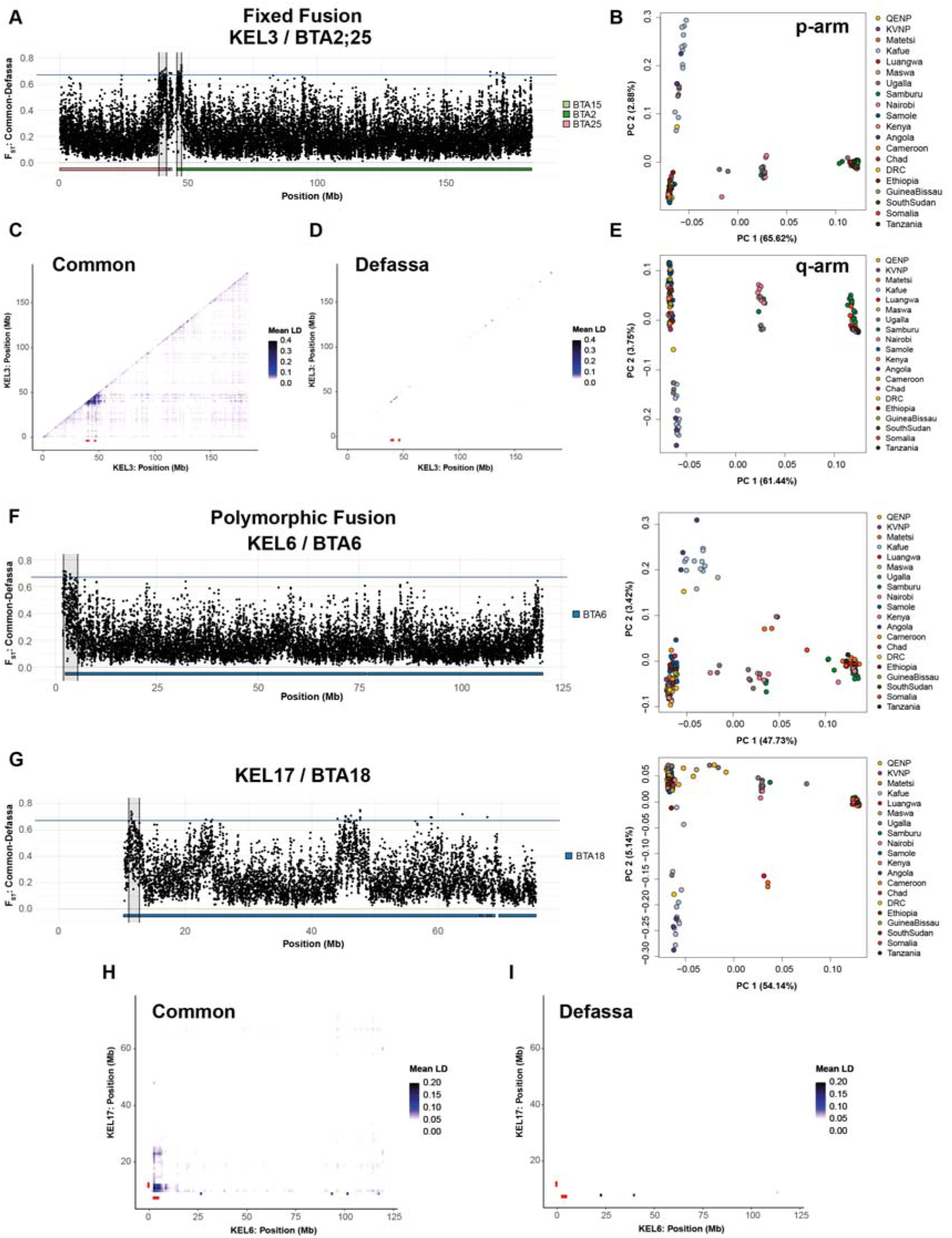
Signatures of Robertsonian (Rb) fusions in waterbuck. (A) F_ST_ of a fixed Rb fusions in KEL3 homologous to cattle BTA2;25. F_ST_ was calculated in 10 Kb windows between subspecies, with the region of interest highlighted in grey and homology to cattle chromosomes plotted below. (B) PCA of the region on the p-arm of KEL3. PCAs were computed for each region using all 143 samples. (C) Mean pairwise LD across KEL3 in common and (D) defassa. Pairwise LD was calculated in 1 Mb windows for each subspecies with regions of interest shown in red. (E) PCA of the region on the q-arm of KEL3. (F) F_ST_ of KEL6 (BTA6) involved in the KEL6;17 (BTA6;18) polymorphic Rb fusion and PCA of the highlighted region. (G) F_ST_ of KEL17 (BTA18) and PCA of the highlighted region. (H) Mean pairwise interchromosomal LD between KEL6 and KEL17 in common and (I) defassa waterbuck.

Our results point to reduced recombination in the pericentromeric region of KEL3, which can lead to increased linked selection, reduced diversity, and increased F_ST_ in these regions. As it has been previously reported, Rbs in both homozygous and heterozygous configurations change the recombination landscape in pericentromeric regions, extending the suppression of recombination further into the interstitial parts of the chromosomes when compared to unfused counterparts (Dumas and Britton-Davidian 2002; Merico et al. 2013), suggesting that our results might be an historical signature of the Rb; however, we cannot discard that the reduction of recombination is only due to the centromeric effect (Talbert and Henikoff 2010; Nambiar and Smith 2016). These results could also be indicative of a novel chromosome rearrangement, with similar indirect methods used in the seaweed fly to detect signatures of putative chromosome inversions (Mérot et al. 2021).

Independently of the mechanism by which reduced recombination in the pericentromeric region of KEL3 happened, we found 128 genes within this genomic region (**Supplementary Table S6**). The region on the p-arm of KEL3 contained several genes involved in male fertility, including the gene IFT140 (F_ST_ = 0.720), associated with ciliated cells such as sperm (Zhang et al. 2018), PRSS21 (F_ST_ = 0.643) involved in proteolytic events in the maturation of testicular germ cells (Netzel-Arnett et al. 2009), and SEP12 (F_ST_ = 0.621) which is involved in the elongation of sperm tails and the morphogenesis of sperm heads, and impacts sperm motility when it becomes phosphorylated (Shen et al. 2017). The region of high F_ST_ on the q-arm contained 15 genes and included OCA2 (F_ST_ = 0.626), part of the mammalian pigmentary system, which has previously been shown to have high F_ST_ between the two subspecies and has been suggested to be involved in the variation in fur colouration within the species (Wang et al. 2024). The gene has been studied in cavefish, where different deletions were found to cause loss of function of the OCA2 protein in two populations, leading to albinism (Protas et al. 2006). Genes in these regions were statistically overrepresented for the GO term pre-synapse (**Supplementary Table S7**).

We then examined the chromosomes involved in the two polymorphic Rb fusions in waterbuck, to investigate the relationship between the fusion event and genomic differentiation and recombination. We found high genomic differentiation (F_ST_) near the centromeres of waterbuck chromosomes KEL6 (BTA6) and KEL17 (BTA18), involved in the KEL6;17 (BTA6;18) Rb fusion (**Fig. 4F and 4G**), with blocks defined between 1,955,000-5,065,000 bp on KEL6 (F_ST_ 99.9^th^ percentile: 0.716) and 11,035,000-12,865,000 bp on KEL17 (F_ST_ 99.9^th^ percentile: 0.735). These regions also had higher intrachromosomal LD in the common subspecies than in the defassa (**Supplementary Fig. S11**). We then calculated interchromosomal LD between KEL6 and KEL17 for each subspecies and found LD was higher in the common waterbuck than the defassa (**Fig. 4H and 4I**). The highest LD was found between regions at the starts of both chromosomes, near the centromeres, and additional regions of moderate interchromosomal LD were found along the chromosomes. The higher interchromosomal LD across the two chromosomes in the common subspecies was expected, as the KEL6;17 Rb fusion is fixed in all karyotypes (2*n* = 50-52) and therefore is a large metacentric Rb chromosome, whilst in the defassa the fusion is either in the heterozygous form (2*n* = 53) or completely absent (2*n* = 54; **Fig 1A**; Kingswood et al., 1998). This suggests the homozygous Rb fusion of KEL6;17 in the common subspecies has resulted in a region of high genomic differentiation and reduced recombination surrounding the centromere, which may also be impacting regions across the Rb fused chromosome, in line with previous observations in mice (Vara et al. 2021; Marín-García et al. 2024). However, we did not find the same signatures for the other polymorphic Rb fusion in waterbuck, KEL8;9 (BTA7;11; **Supplementary Fig. S13**). This could be because the high F_ST_ in the pericentromeric region in the polymorphic Rb KEL6;17 is not a hallmark of the Rb, or because KEL8;9 is only present in in the 2*n* = 51 (heterozygous) and 2*n* = 50 (homozygous) karyotypes in the common subspecies (**Fig. 1A**; Kingswood et al., 1998), and these are either less frequent in wild populations and/or under sampled in our genomic study.

PCAs of the two regions near the centromeres of KEL6 and KEL17 resulted in three groupings on PC1 (with a variance of 47.73% and 54.1.4%; **Fig. 4**), supporting a polymorphic variation. The left group contained individuals from defassa waterbuck populations (except the highly admixed individual from Nairobi population, Fig. 2c), the right group contained exclusively individuals of the common waterbuck populations, while individuals in the centre group belonged to populations along the contact zone for both subspecies (common: Luangwa, Matetsi, Nairobi and Samburu, and defassa: Ugalla). Following previous publications (e.g., Mérot 2020), local PCA clusters could reflect the three haplotypes of the Rb fusion (wildtype, heterozygous, and homozygous), however this would suggest that the common subspecies is also polymorphic for the KEL6;17 fusion, having the heterozygous fusion, which has not been documented in previous cytogenetic studies (Kingswood et al. 1998; Pagacova et al. 2011). These studies have focused predominantly on captive samples, whereas our study involves wild samples from across most of the waterbuck’s distribution and therefore these karyotypes may have previously been unsampled. Further captive sampling of waterbuck karyotypes in Pagacova et al. 2011 found evidence of another Rb fusion in the defassa subspecies (BTA7;29), suggesting that other combinations of Rb fusions are possible in this species.

On KEL6 a total of 16 annotated genes were found in the region of interest, whilst 42 genes were found on KEL17 (**Supplementary Table S6**). These two regions included several genes such as APELA (F_ST_ = 0.565) which is involved in early embryogenesis (Chng et al. 2013; Pauli et al. 2014; Norris et al. 2017) and the gene TERF2IP/RAP1 (F_ST_ = 0.532) that is associated with the shelterin complex, protecting telomeres from DNA damage and fusion events (De Lange 2005). RAP1 regulates telomere recombination by homologous directed repair, increasing fragility (Martinez et al. 2010). Telomere loss or inactivation is required for the formation of Rb fusions (Slijepcevic 1998) and therefore this gene may play a role in the formation of Robertsonian fusions. A total of 12 GO terms were statistically overrepresented in the F_ST_ block on KEL6 and one on KEL17 (**Supplementary Table S7**).

These results provide support for the recombination suppression model of chromosome evolution (Rieseberg 2001; Farré et al. 2013). We find regions of high LD surrounding the centromeres of some chromosomes involved in fixed and polymorphic Rb fusions, indicating reduced recombination within these areas. This has led to high levels of genomic differentiation and local population structure, potentially linked to the different karyotypes. Moreover, we find genes within these regions that are involved in fertility, development, and recombination, key processes in reproductive isolation.

## Conclusion

In conclusion, we provide new insights into population dynamics and chromosome evolution in waterbuck. Through the assembly of a high-quality chromosome-level genome and re-sequencing of historical museum collections, we were able to uncover fine-scale population structure and gene flow across most of the species’ distribution in Africa. These findings indicate that population structure largely reflects geographic locality, with the Great Rift Valley and the Congolian Rainforest likely serving as barriers to gene flow between subspecies and north/south groups.

Across waterbuck chromosomes, we identified several regions exhibiting high levels of genomic differentiation, with some of these near the centromeres of chromosomes involved in Rb fusions. Here, we provide putative evidence of Rb fusions impacting genomic differentiation, recombination, and population structure within the waterbuck. These regions also contained genes involved in fertility, development, and recombination. However, the role of Rbs in conferring reproductive isolation in waterbuck is still debatable. We found admixed individuals in the hybrid zone (**Fig. 2C**) with potentially heterozygous karyotypes (**Fig. 4F and 4G**), suggesting that if any, these Rbs might not provide a strong genetic barrier. Karyotyping the animals used in this study, for example using Hi-C sequencing, is needed to further elucidate the role of Rbs in waterbuck evolution. The species is therefore an interesting model to further explore intraspecies chromosome evolution.

Our work also has important applications for conservation. Historical sampling has improved our understanding of population structure within this species, and our comprehensive dataset provides a framework for future management strategies. From these results it is clear that each subspecies and north/south group should be managed separately, with careful consideration of populations along the entire contact zone where admixture has been found. Finally, our study underscores the importance of considering chromosome rearrangements in conservation efforts and highlights the need for improved methods to karyotype wild animals.

## Methods

### Primary mammalian cell culture and karyotyping

A primary fibroblast mammalian cell line was established for a captive female defassa waterbuck (*Kobus ellipsiprymnus defassa*) from a tissue sample provided by the Aspinall Foundation (Kent, UK). The culture was maintained in DMEM containing 10% FBS and 1% Pen-Strep, incubated at 37°C with 5% CO_2_. Cells were harvested on several passages and the stability of the karyotype was checked. Harvesting of chromosomes followed (Howe et al. 2014). Briefly, colcemid (10 µl/ml) was added to cell culture flasks and incubated at 37°C with 5% CO_2_ for 45 min. Cells were washed with 1x PBS. Trypsin was added to the flasks for 2 min before transferring contents to a tube. Tubes were centrifuged at 200 x g for 5 min, followed by removal of the supernatant and resuspension in 5 ml of Carnoy’s Fixative (3:1 methanol and glacial acetic acid) with vortexing. A further 5 ml of fixative was added without vortexing, centrifuged at 200 x g for 5 min, and the supernatant discarded. This step was then repeated, and chromosome preps were stored at 4°C. Slides were stained with DAPI, and the metaphases were captured with fluorescent microscopy and karyotyped using SmartTypeDemo v3.3.2.

### DNA extraction and long-read PacBio HiFi sequencing

DNA was extracted from the primary cell line using the QIAGEN Blood and Cell DNA extraction kit following the cell culture protocol, including an RNase A treatment before lysis. DNA was eluted in 10 mM Tris-HCl (pH 8.5) and quantified using NanoDrop and Qubit. DNA was also measured for fragment lengths using pulse field gel electrophoresis (PFGE). DNA was sent to Edinburgh Genomics (Edinburgh, UK), where one PacBio SMRTbell library was prepared and sequenced on two PacBio Sequel IIe SMRT 8M Cells in HiFi mode.

### De novo genome assembly and gene annotation

The pipeline for genome assembly was adapted from the Vertebrate Genome Project (VGP) pipeline and is summarised below. HiFi reads in BAM format were converted to FASTQ format using Samtools v1.6 ‘fastq’ (Danecek et al. 2021), followed by quality control with FASTQC v0.11.9 (https://www.bioinformatics.babraham.ac.uk/projects/fastqc), MultiQC v1.0.dev0 (Ewels et al. 2016), and Nanoplot v1.32.1 (De Coster et al. 2018). Reads containing the adapters (ATCTCTCTCAACAACAACAACGGAGGAGGAGGAAAAGAGAGAGAT and ATCTCTCTCTTTTCCTCCTCCTCCGTTGTTGTTGTTGAGAGAGAT) were removed with Cutadapt v1.18 (Martin 2011) with parameters minimum error rate 0.1%, minimum overlap length 35, minimum length 5000 bp, maximum length 30000 bp, and discard trimmed reads. To estimate the genome size, the maximum read depth, and the transition coverage between the haploid and diploid peaks we used a k-mer-based approach (32-mer) with Meryl v1.3 (Rhie et al. 2020) and profiled using GenomeScope2 v2.0 (Ranallo-Benavidez et al. 2020). The genome was assembled with Hifiasm v0.16.1-r375 (Cheng et al. 2021), with purging of all types of haplotigs, a purge maximum of 45, and removal of 20 bp from both ends of the reads, which output a primary contig-level genome assembly. Additionally, the mitochondrial genome was assembled with the MitoHiFi v2.2 pipeline (Uliano-Silva et al. 2023). Genome statistics were produced with QUAST v5.0.2 (Mikheenko et al. 2018) and Merqury v1.3 (Rhie et al. 2020), and the genome was assessed for completeness and contamination with the BlobToolKit v2 pipeline (Challis et al. 2020) using BUSCO v5 (Manni, Berkeley, Seppey, and Zdobnov 2021; Manni, Berkeley, Seppey, Simão, et al. 2021), BLAST v2.10.0, and Uniprot 2023_01 (Consortium 2023) databases.

The Hi-C library was prepared from the defassa waterbuck primary cell line and sequenced as described in Álvarez-González et al., 2022. Additional Hi-C data was generated by DNA Zoo (https://www.dnazoo.org) following the protocol in Rao et al. 2014 from a female common waterbuck blood sample bred in captivity, with a karyotype of 2*n* = 52. Both Hi-C datasets were aligned to the contig-level genome assembly with Juicer (Durand et al. 2016), however due to high levels of interchromosomal Hi-C contacts in the defassa Hi-C dataset, the common waterbuck Hi-C dataset was used to scaffold the genome using 3D-DNA (Dudchenko et al. 2017).

The chromosome-level genome was reviewed and curated using Juicebox Assembly Tools (JBAT; Dudchenko et al., 2018). The defassa Hi-C dataset was used to resolve the karyotype to 2*n* = 54 and this was confirmed via synteny to the cattle reference genome (*Bos taurus*; ARS-UCD2.0) and the previous cytogenetic study (Kingswood et al. 1998). The waterbuck genome was aligned to cattle with MashMap (Jain et al. 2018), syntenic blocks were constructed at 300 Kb resolution using several customised scripts, and then visualised with syntenyPlotteR v1.0.0 (Quigley et al. 2023) in R. Scaffolded chromosomes orthologous to BTA6 and BTA18 in the draft assembly were split into two chromosomes in the chromosome-level waterbuck genome assembly. An interactive map was produced showing two Hi-C datasets produced by DNA Zoo (2*n* = 52 and an additional unstudied karyotype; 2*n* = 51) aligned to the chromosome-level genome assembly (https://www.dnazoo.org/post/don-t-go-chasing-waterbuck).

In this study, we further curated the genome assembly by reorientating chromosomes with their centromeres closer to the start and reordering by size (**Supplementary Table S1**). Both sets of Hi-C data (2*n* = 52 and 2*n* = 54) were then remapped to this final curated assembly, where the common Hi-C data had strong interchromosomal interactions between waterbuck chromosomes KEL6 and KEL17, syntenic to BTA6 and BTA18, respectively (**Fig. 1D and Supplementary Fig. S3B**), whilst the defassa Hi-C did not have stronger interchromosomal interactions compared to other chromosomes (**Supplementary Fig. S3C**). The chromosome-level genome assembly was annotated for repeats with RepeatMasker v2.6.0+ (http://www.repeatmasker.org) using the Dfam_Consensus-20181026 and RepBase-20181026 databases, and the *Bos taurus* dataset. Protein-coding genes were annotated with GeMoMa v1.9 (Keilwagen et al. 2016; Keilwagen et al. 2018) using both goat (*Capra hircus*; ARS1.2) and cattle (*Bos taurus*; ARS-UCD2.0) as reference. Genes were filtered to exclude duplicate gene annotations with multiple transcripts or genes that had multiple annotations but the same coordinates.

### Circos plot

To visualise the final genome assembly and annotation, 100 Kb windows were created along each of the 27 chromosomes. The percentage of GC content was calculated with SeqKit v2.6.1 fx2tab (Shen et al. 2016). Repeat density was calculated as the total number of masked bases in each window divided by 100 Kb. Gene density was calculated as the number of protein-coding genes in each 100 Kb window. Visualisation was performed using Circos 0.69.8 (Krzywinski et al. 2009).

### Historical museum sampling and sequencing

Historical waterbuck skin samples were provided by the Powell Cotton Museum (Kent, UK) and the Royal Museum for Central Africa (Tervuren, Belgium). DNA was extracted from skin samples using a modified phenol-chloroform protocol, adapted from McDonough et al., 2018, Roycroft et al., 2021 and Molecular Cloning Vol I, and described as follows. Skin samples were dissected into small sections (5-10 mm) using sterile equipment, briefly cleaned in a dilute bleach solution, cleaned with sterile ultrapure water. Samples were then incubated in water at room temperature, with frequent water changes, to clean and rehydrate the skin. Water was removed after 48 hours and 320 μl lysis buffer (100 mM Tris-HCl pH 8.0, 5 mM EDTA pH 8.0, 200 mM NaCl, and 0.2% SDS; Z. Wang & Storm, 2006), 40 μl Proteinase K, and 40 μl DTT were added to each tube. Samples were incubated at 56°C for up to 48 hours and vortexed regularly. DNA was extracted using a standard phenol-chloroform protocol with QIAGEN MaXtract tubes and eluted in TE buffer (pH 8.5). DNA was measured for quantity using NanoDrop and Qubit, and DNA length using gel electrophoresis. DNA extraction and quality control took place in a room dedicated to pre-PCR work. A total of 24 samples were sent for Illumina library preparation (without size selection) and whole genome sequencing (WGS) at Novogene (UK), with a 2 x 150 bp read length and 350 bp insert size, on an Illumina NovaSeq6000.

### Mapping of WGS data and filtering

Historical WGS data was quality controlled with FASTQC v0.11.9 and MultiQC v1.0.dev0 (Ewels et al. 2016) and the adapters (AGATCGGAAGAGCGTCGTGTAGGGAAAGAGTGTAGATCTCGGTGGTCGCCGTATCATT and GATCGGAAGAGCACACGTCTGAACTCCAGTCACGGATGACTATCTCGTATGCCGTCTTCTGCTTG) were trimmed and reads collapsed using AdapterRemoval v2.3.3 (Schubert et al. 2016) with parameters --mm 3, --collapse, --collapse-conservatively, --trimns, and --trimqualities. Paired-end and collapsed reads were aligned separately with BWA-MEM v0.7.17 (Li 2013) to the assembled chromosome-level waterbuck genome. BAM files were fixed and checked with Picard v3.0.0 FixMateInfomation and ValidateSAMFile (https://www.broadinstitute.github.io/picard/), and Samtools v1.6 calmd (Danecek et al. 2021). Duplicates were removed from paired-end and collapsed BAM files with Picard v3.0.0 MarkDuplicates and the PALEOMIX v1.3.7 rmdup_collapsed script (Schubert et al. 2014), respectively. DNA damage was assessed, and quality scores were rescaled based on damage, using mapDamage v2.2.0 (Jónsson et al. 2013). Paired-end and collapsed BAM file flags were modified with Samtools v1.6 view and merged with Samtools v1.6 merge (Danecek et al. 2021). Mapping statistics for the merged alignment files were calculated with Samtools v1.6 stats and depth (Danecek et al. 2021) and Qualimap v2.2.2-dev (Okonechnikov et al. 2016). Moreover, published WGS reads from 119 modern waterbuck samples from 10 populations (Wang et al. 2024) was mapped to the chromosome-level reference genome using a modified version of the PALEOMIX bam pipeline (Schubert et al. 2014). Lastly, BAM files were filtered for secondary and supplementary alignments, PCR duplicates, quality, insert sizes less than 50lllbp or greater than 1,000lllbp, reads with less than 50lllbp or 50% mapped, and reads mapping to different scaffolds or with unexpected orientations.

Genomic sites were then filtered in both the reference genome and the alignment files as described in (Wang et al. 2024). Briefly, genomic sites on unplaced scaffolds and sex chromosomes, and sites annotated as repetitive, were firstly removed. Then regions with excess heterozygosity were identified in the 24 historical BAM files and removed. The historical BAM files were also used to calculate the global depth across WGS samples and remove sites below the 1^st^ percentile and above the 99^th^ percentile. Finally, sites with low mappability were excluded. This resulted in a filtered genomic sites file which was used when calling genotype likelihoods.

### Population structure

Genotypes likelihoods were called per chromosome on filtered sites with ANGSD v0.940 (Korneliussen et al., 2014; -GL 2, -doMajorMinor 1, -doMaf 1, -SNP_pval 1e-6, -minMaf 0.05, -minMapQ 30, and -minQ 20). Genotypes were also calculated for only transversion sites (-rmtrans 1). Beagle likelihood files were combined for all chromosomes and samples, and the combined beagle file was used in both the principal component analyses (PCA) and admixture analyses. PCAs were constructed using pcangsd v.0.99 (Meisner and Albrechtsen 2018) with -minMaf 0.05, whilst admixture proportions were calculated with ANGSD v0.940 NGSadmix (Skotte et al. 2013) with -minMaf 0.05 and estimated populations of K between K = 2 and K = 20, with K = 2, K = 3, K = 4, and K = 12 plotted in this study using R as they were the most informative. NJ trees were constructed with ANGSD v0.940 (Korneliussen et al., 2014; -GL 2, -minMapQ 30, -minQ 20, -doMajorMinor 1, -doMaf 1, -SNP_pval 1e-6, -doIBS 1, -doCounts 1, -doCov 1, -makeMatrix 1, and -minMaf 0.05). The NJ tree was constructed using the pairwise IBS matrix with a custom R script.

Estimated Effective Migration Surfaces (EEMS) were calculated by firstly running ANGSD v0.940 (Korneliussen et al. 2014) using parameters -GL 2, -minMapQ 30, -minQ 20, -doMajorMinor 1, -doMaf 1, -SNP_pval 1e-6, -doIBS 1, -doCounts 1, -doCov 1, -makeMatrix 1, and -minMaf 0.05. The EEMS runeems_snps pipeline (Petkova et al. 2015) was run in triplicates (nIndiv 143, nSites 143, nDemes 600, diploid, numMCMCIter 20000000, numBurnIter 10000000, and numThinIter 9999) and visualised with the library rEEMSplots in R.

### Heterozygosity

To calculate overall genome-wide heterozygosity per sample for filtered sites, ANGSD v0.940 (Korneliussen et al. 2014) was used (-doSaf 1, -GL 2, -minMapQ 30, and -minQ 20) to produce site allele frequency (SAF) files, with the reference genome used as the ancestral state. Transversions were also removed (-noTrans 1) to consider differences between all sites and transversion-specific sites. SAF files were then converted into the folded site frequency spectrum (SFS) files by the ANGSD v0.940 realSFS program (Korneliussen et al. 2014) and plotted in R.

### Genomic differentiation and detecting signatures of putative chromosome rearrangements

Genomic differentiation (F_ST_) was estimated between the two subspecies of waterbuck, *Kobus ellipsiprymnus ellipsiprymnus*(common) and *Kobus ellipsiprymnus defassa* (defassa), by running ANGSD v0.940 (Korneliussen et al. 2014) separately for the two subspecies on filtered sites (-GL 2, -doSaf 1, -minMapQ 30, and -minQ 20). Folded SFS files were generated with ANGSD v0.940 realSFS (Korneliussen et al. 2014) using the waterbuck reference genome as the ancestral state (-fold 1), followed by fst index (-whichFst 1 and -fold) and fst stats2 (window size and step size of 10 Kb). To find genes within the F_ST_ windows across the genome, the output file was intersected with the waterbuck homology-based gene annotation file using BEDTools intersect v2.31.0 (Quinlan and Hall 2010). Plots were created from F_ST_ values with the addition of synteny data to cattle chromosomes using a custom R script.

LD was calculated using several methods. Genotype likelihoods were computed for each chromosome on filtered genomic sites with ANGSD v0.940 (Korneliussen et al. 2014) using parameters -minMapQ 30, -minQ 20, -doCounts 1, -GL 2, -doMajorMinor 1, -doMaf 1, -skipTriallelic 1 -doGlf, -SNP_pval 1e-6, -remove_bads, -minMaf 0.05. The program ngsLD v1.2.1 (Fox et al. 2019) was used with parameters --max_kb_dist 1000 and --rnd_sample 0.1. Mean LD was then calculated in 100 Kb windows across chromosomes using a custom R script. Mean pairwise LD was also calculated in 1 Mb windows on chromosomes involved in putative rearrangements using a custom R script. We also calculated interchromosomal LD between KEL6 and KEL17 by computing genotype likelihoods on the two chromosomes and calculating LD using ngsLD (Fox et al. 2019) with the parameters --max_kb_distance 0 and --rnd_sample 0.001. Files were filtered for interchromosomal interactions and mean pairwise LD was calculated in 1 Mb windows.

Local PCAs of particular regions of interest were computed by filtering the genomic sites file and genotyping with ANGSD v0.940 (Korneliussen et al. 2014). The PCAs were then computed on the 143 individuals and visualised as described previously.

Gene ontology (GO) statistical overrepresentation tests were computed with Panther (Mi et al. 2019; Thomas et al. 2022) with *Bos taurus* as reference on lists of waterbuck genes found within the 99^th^percentile of F_ST_ windows or in blocks of high F_ST_.

## Acknowledgements

We thank the specialist High Performance Computing system provided by Information Services at the University of Kent, the Powell Cotton Museum and the Royal Museum for Central Africa for access to their museum collections, and Port Lympne Reserve and The Aspinall Foundation for providing captive samples. The Hi-C guided assembly was done in association with the DNA Zoo consortium (dnazoo.org), which acknowledges support from Illumina, IBM, and Pawsey Supercomputing Center. We thank Prof Terry Robinson for insightful discussions on previous versions of the manuscript.

C.K. was supported by a PhD scholarship from the University of Kent’s Global Challenges Doctoral Centre. M.F. is supported by the Royal Society Research Grant (RGS\R1\211047). C.C.-R. was supported by the GTA fellowship programme from the University of Kent. A.R-H is supported by the Spanish Ministry of Science and Innovation (PID2020–112557GB-I00 to AR-H), the Agència de Gestió d’Ajuts Universitaris i de Recerca, AGAUR (2021 SGR 0122 to A.R-H), and the Catalan Institution for Research and Advanced Studies (ICREA). L.A.-G. was supported by a FPI predoctoral fellowship from the Ministry of Economy and Competitiveness (PRE-2018-083257). E.L.A. acknowledges support from the Behavioral Plasticity Research Institute (NSF DBI-2021795), NSF Physics Frontiers Center (NSF PHY-2210291), and an NIH CEGS (RM1HG011016-01A1) award.

## Supplementary Data

The chromosome-level genome prior to reordering and reorientation of chromosomes can be found at DNA Zoo (https://www.dnazoo.org/assemblies/kobus_ellipsiprymnus) and NCBI SRA (BioProject PRJNA512907). The historical WGS data can be found at NCBI SRA (BioProject PRJNA1232813). Custom code used as part of this project can be found at https://www.github.com/Farre-lab/Kirkland_Waterbuck.

