## Supplementary Figures for "Chromosome-level genomics and historical museum collections reveal new insights into the population structure and chromosome evolution of waterbuck"

### Supplementary Materials

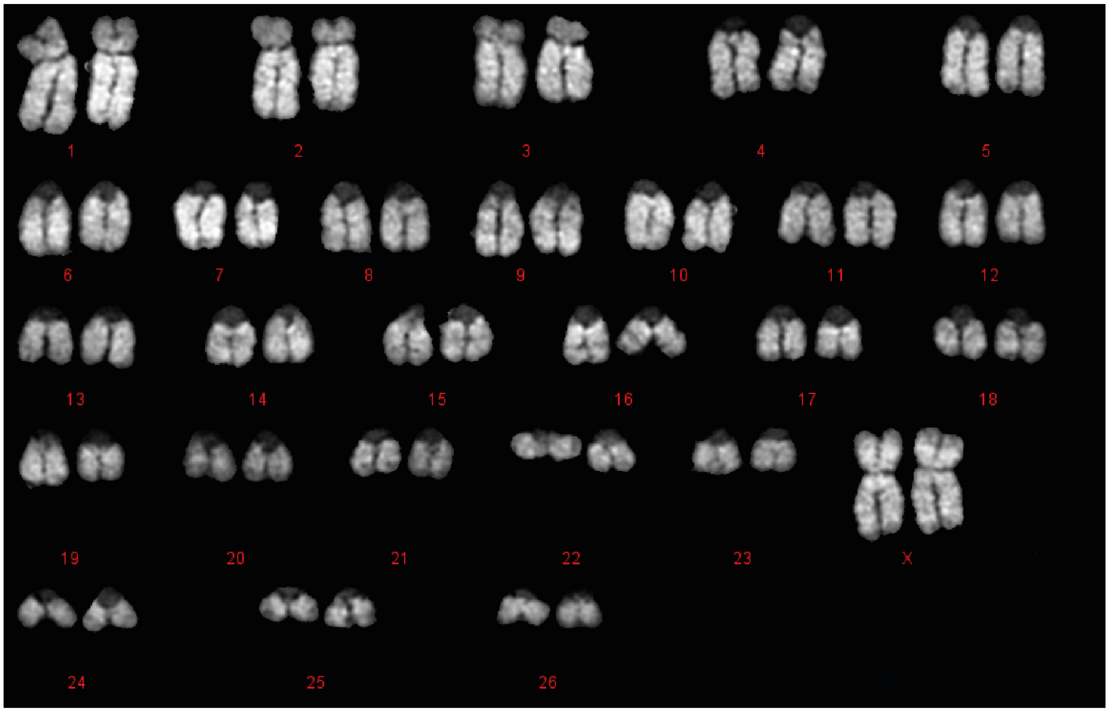


**Supplementary Fig. S1:** Defassa waterbuck cell culture karyotype (2*n* = 54, XX) stained with DAPI, with chromosomes named by size and the X chromosomes labelled separately.

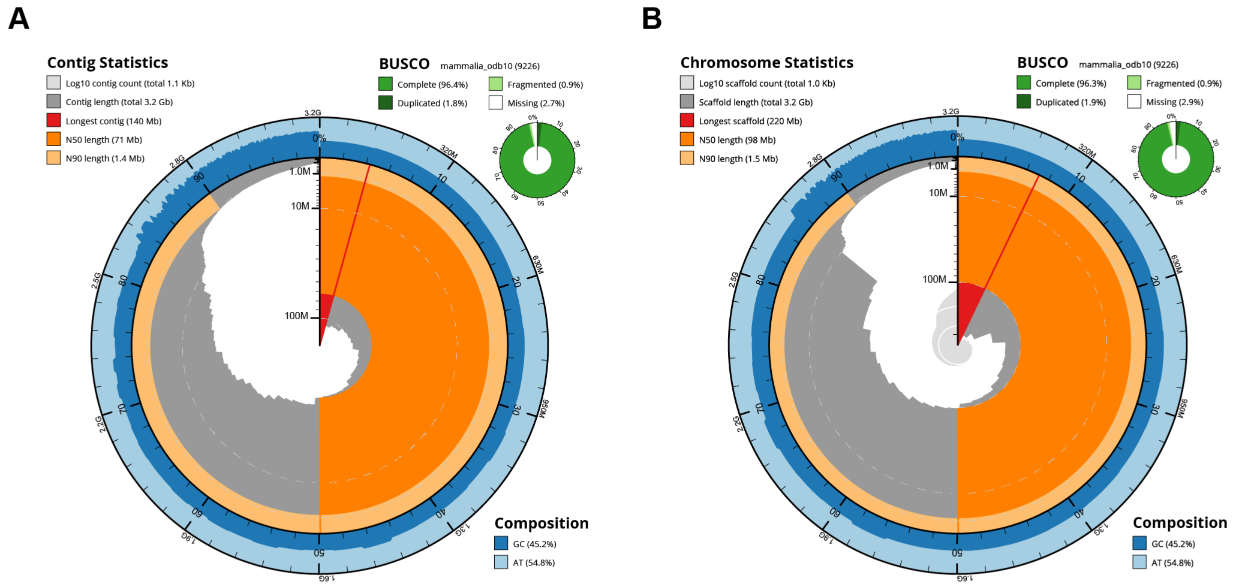


**Supplementary Fig. S2:** Snail plot of the Defassa waterbuck (A) contig-level and (B) chromosome-level genome assemblies.

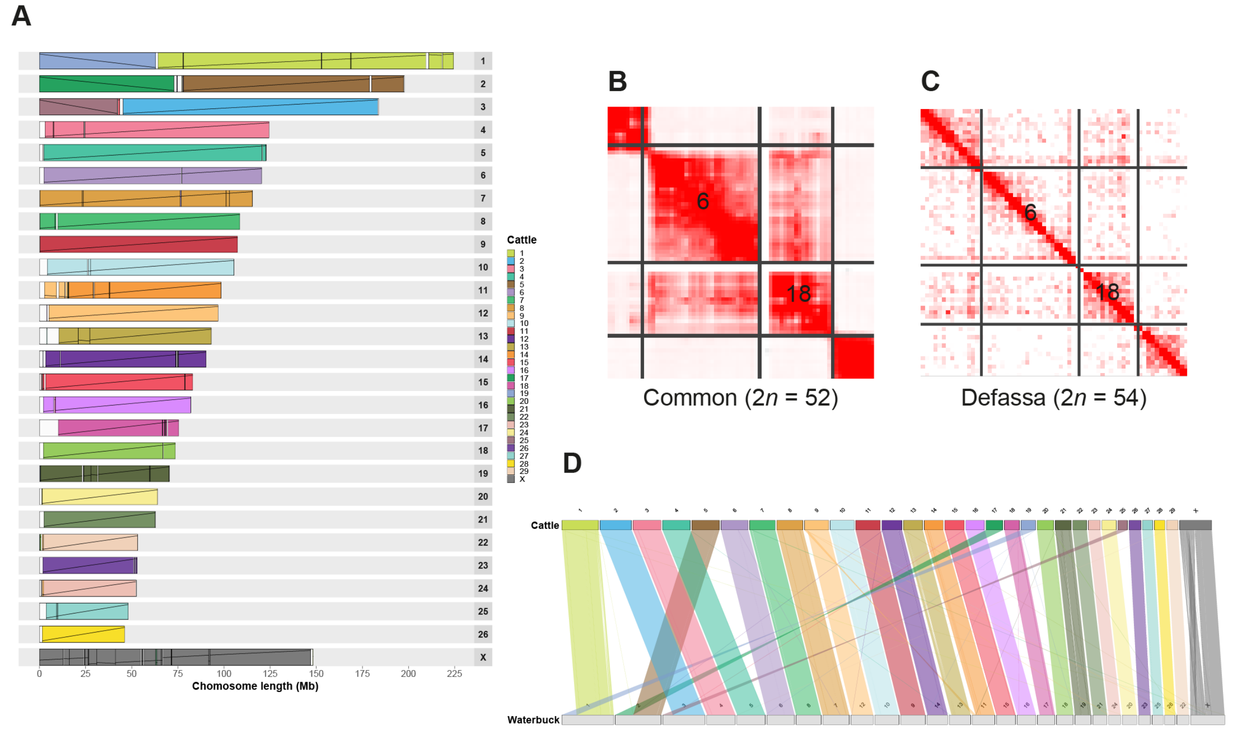

**Supplementary Fig. S3:** Waterbuck chromosome-level assembly after final genome curation. (A) Chromosome painting plot of the synteny between waterbuck and cattle genomes, with colours representing homology to cattle chromosomes. (B) Hi-C interaction matrix of the Hi-C sample 2*n* = 52, showing interactions between KEL6 and KEL17 (homologous to cattle BTA6 and BTA18, respectively) and (C) Hi-C sample 2*n* = 54. (D) Linear plot of the synteny between waterbuck and cattle chromosomes.

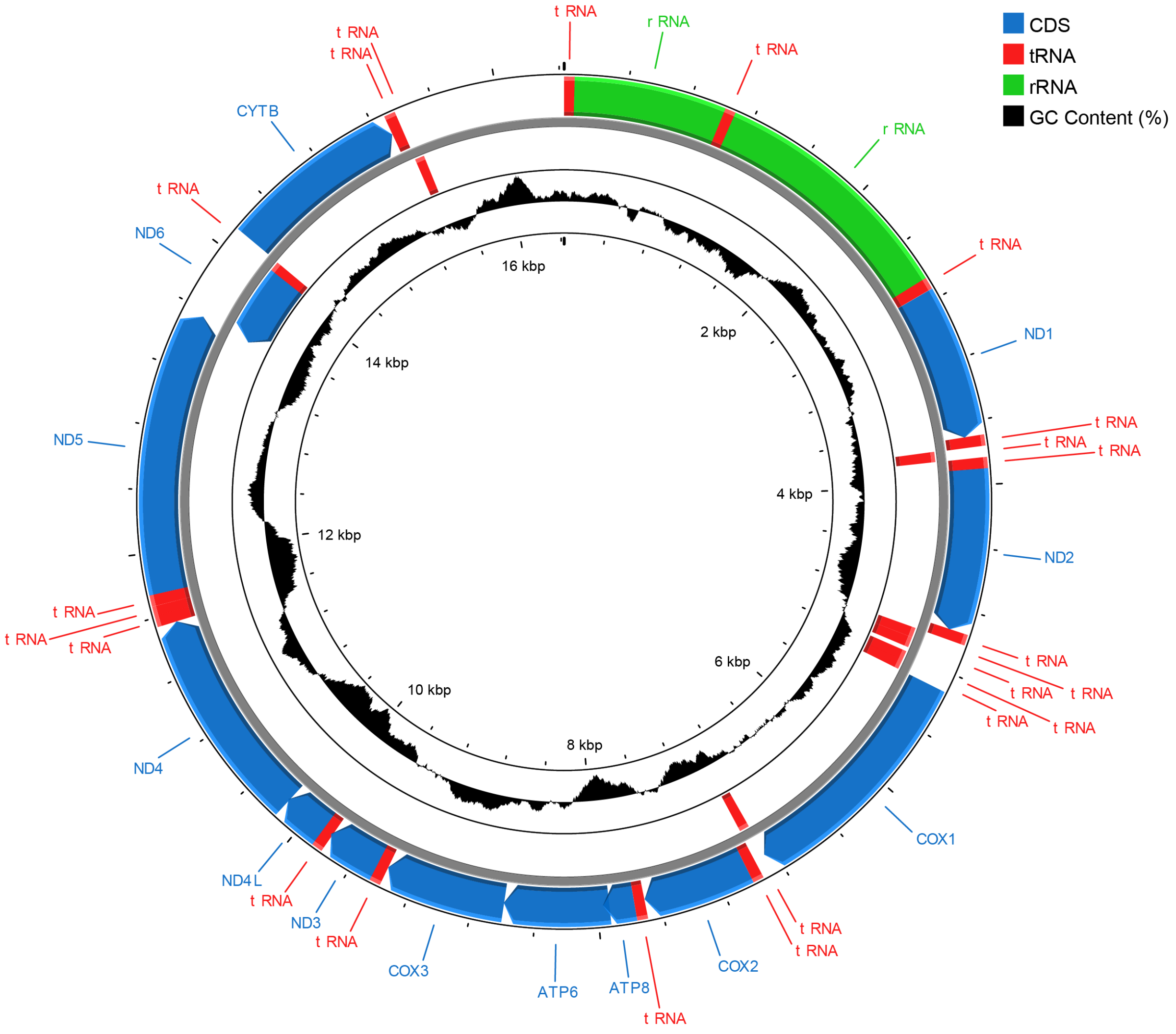


**Supplementary Fig. S4:** Mitochondrial genome assembled from the Defassa waterbuck PacBio HiFi reads, with coding sequences (CDS), tRNAs, rRNAs, and GC content (%) annotated.

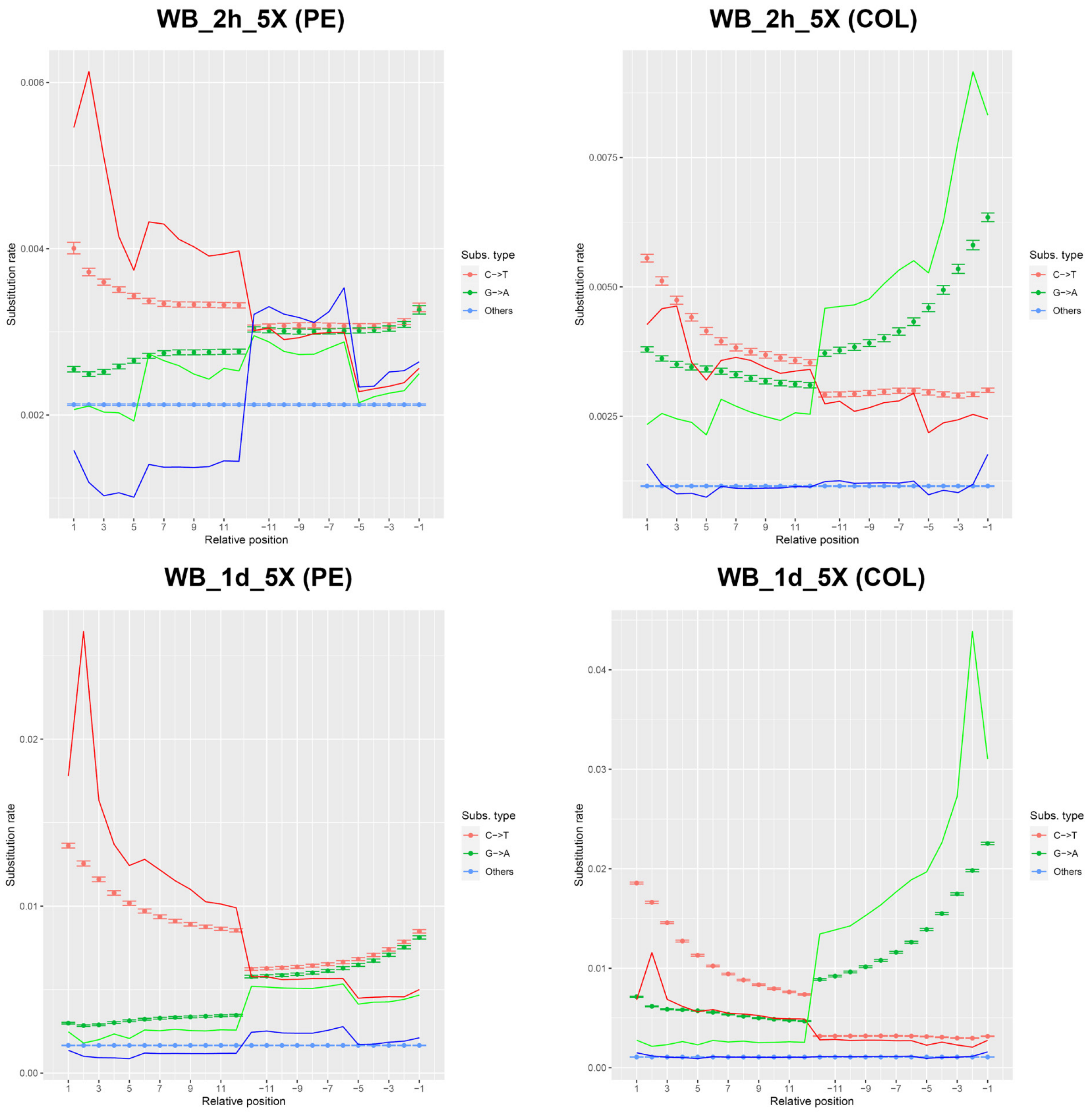


**Supplementary Fig. S5:** Estimation of DNA damage in the historical samples. Two of the samples with the highest (WB_1d_5X) and lowest (WB_2h_5X) substitution rates are shown. Both paired-end (PE) and collapsed (COL) mapped reads were analysed for each sample. Relative position refers to the position at the start and end of the read. Empirical misincorporation frequencies are denoted by the soil line. 95% simulated posterior predictive intervals of the fitted model are shown by the confidence intervals.

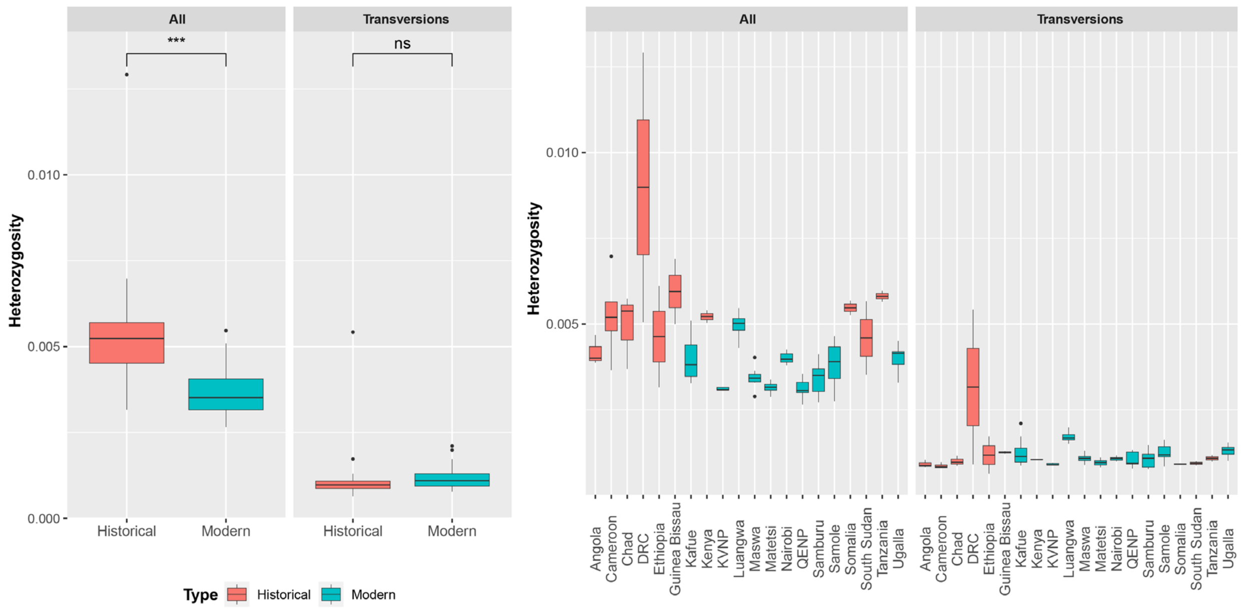


**Supplementary Fig. S6:** Genome-wide heterozygosity calculated for each sample using either all or transversion genomic sites. Samples are grouped by type (historical or modern), or by population.

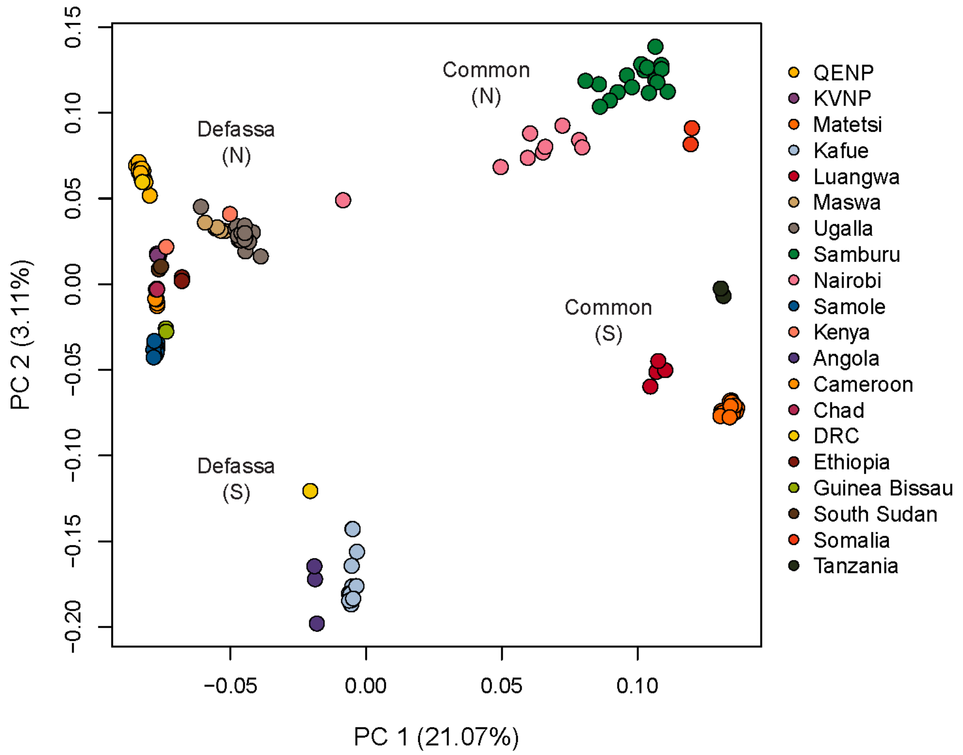


**Supplementary Fig. S7:** PCA of the 143 waterbuck samples using transversion genomic sites and coloured by population.

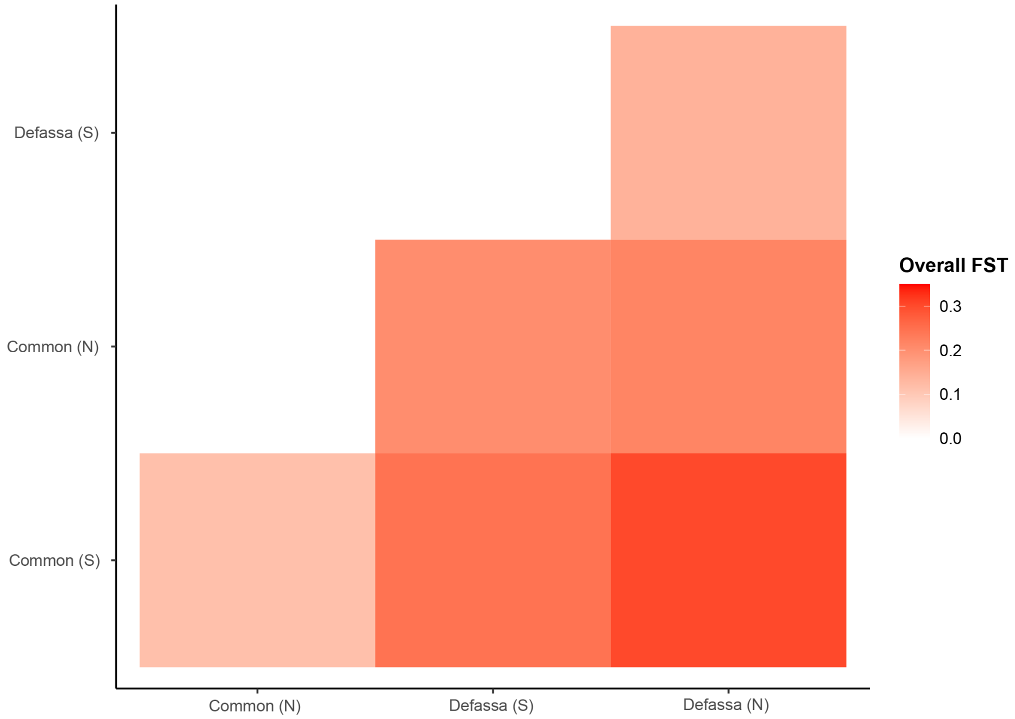


**Supplementary Fig. S8:** Overall $F_{ST}$ between the waterbuck groups.

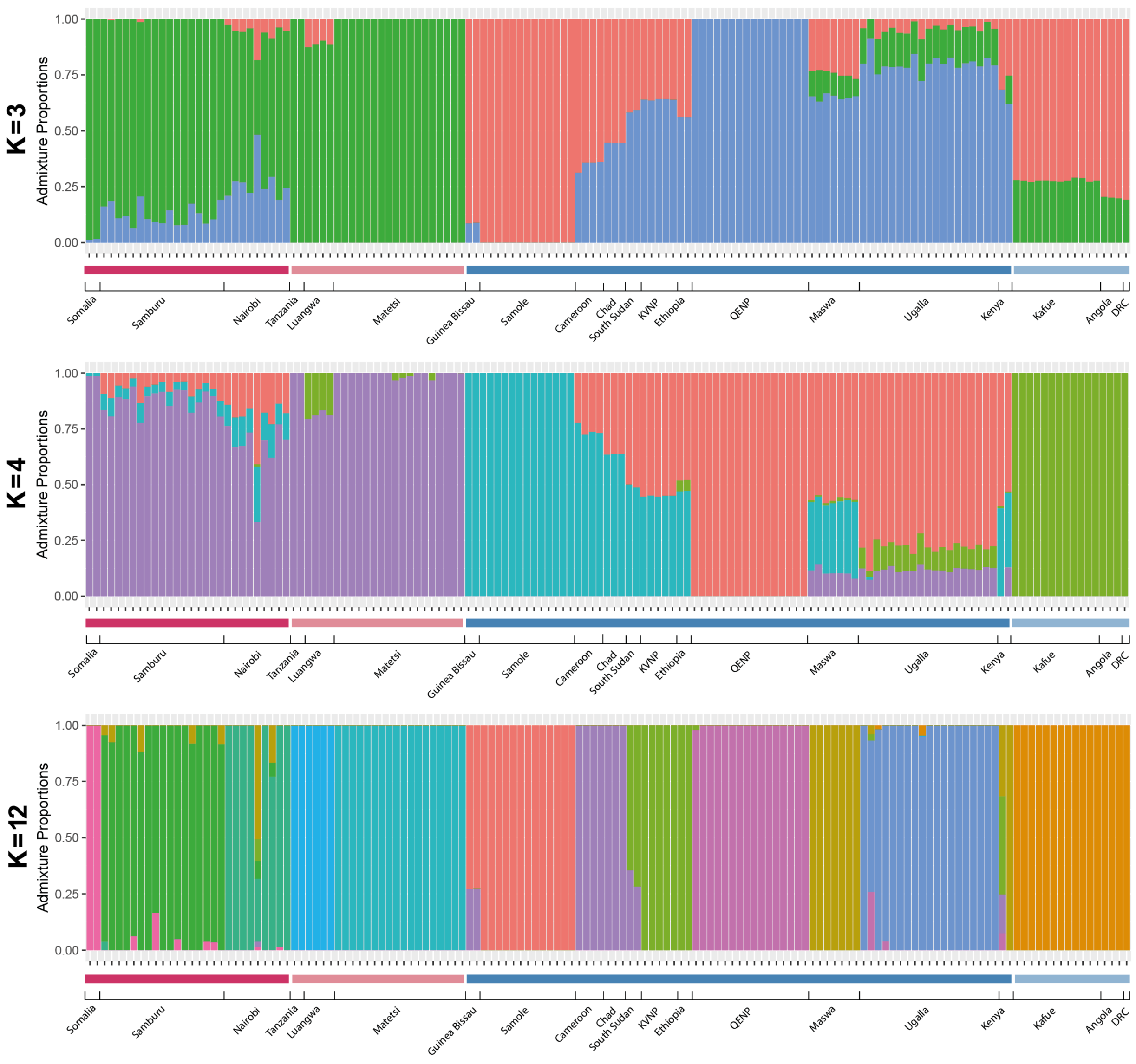


**Supplementary Fig. S9:** Admixture proportions of the waterbuck samples at K = 3, K = 4, and K = 12 estimated populations. Dark red is common (N), light red is common (S), dark blue is Defassa (N), and light blue is Defassa (S).

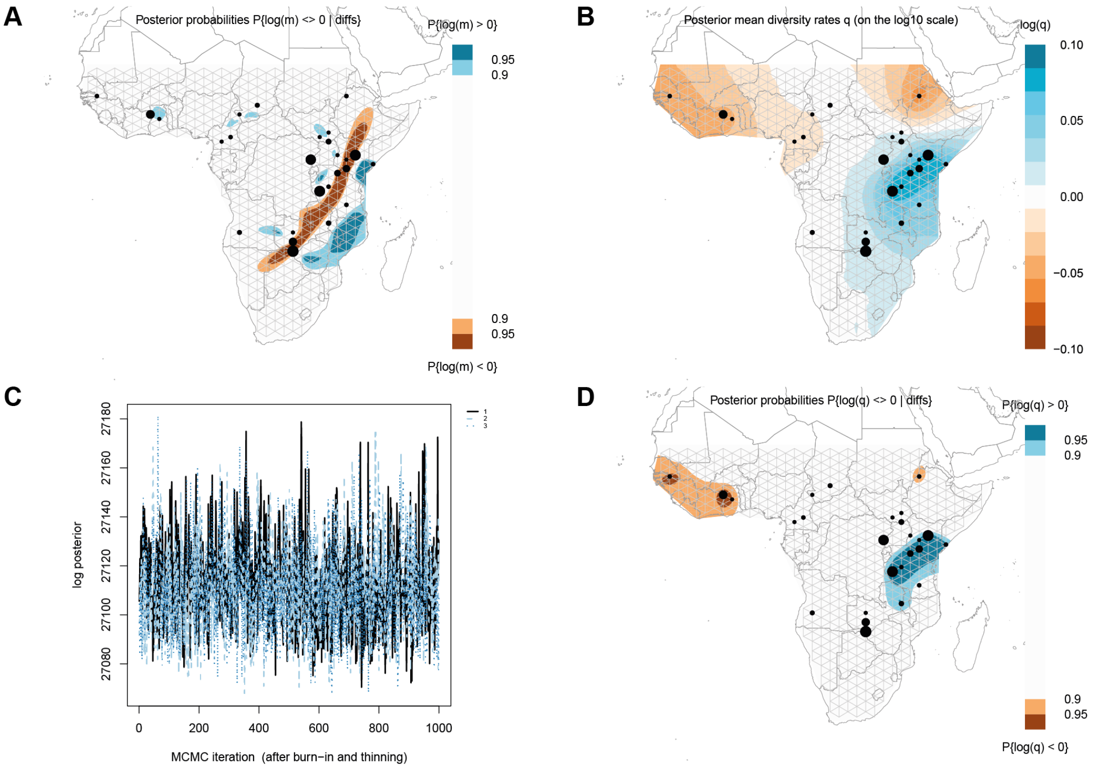


**Supplementary Fig. S10:** EEMS analysis of 142 waterbuck samples, excluding “WB_3k_5X”. (A) Maximum posterior probabilities of the migration rate (*m*). (B) Posterior mean diversity rates (*q*). (C) MCMC iterations showing convergence. (D) Maximum posterior probabilities of the diversity rate. Populations denoted by black circles, with size proportional to the number of individuals in each population.

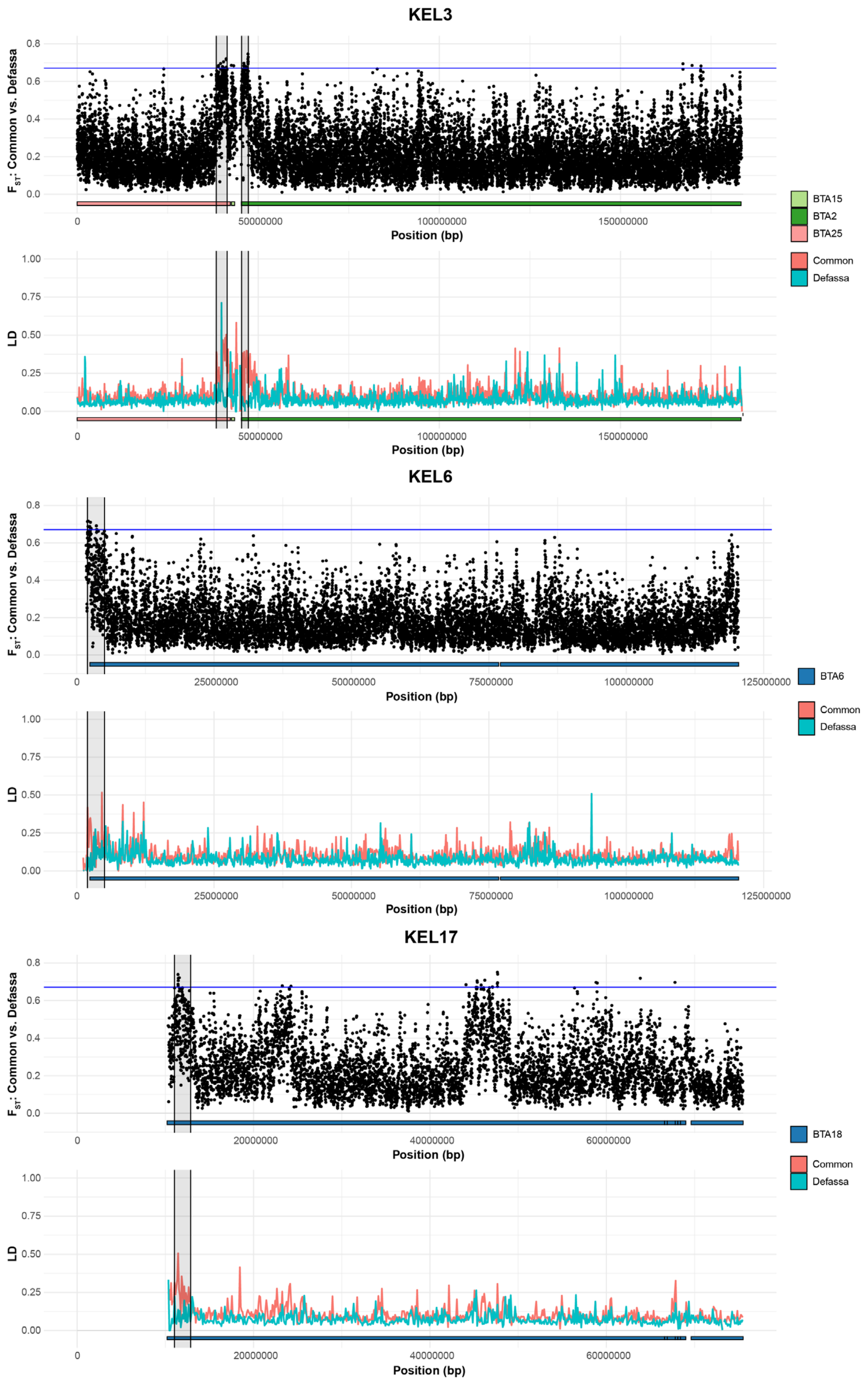


**Supplementary Fig. S11:** $F_{ST}$ between waterbuck subspecies and mean LD in 100 Kb windows for each subspecies across chromosomes KEL3, KEL6, and KEL17 with fixed or polymorphic Rb fusions. Homology to cattle chromosomes (BTA) shown. Regions of interest highlighted.

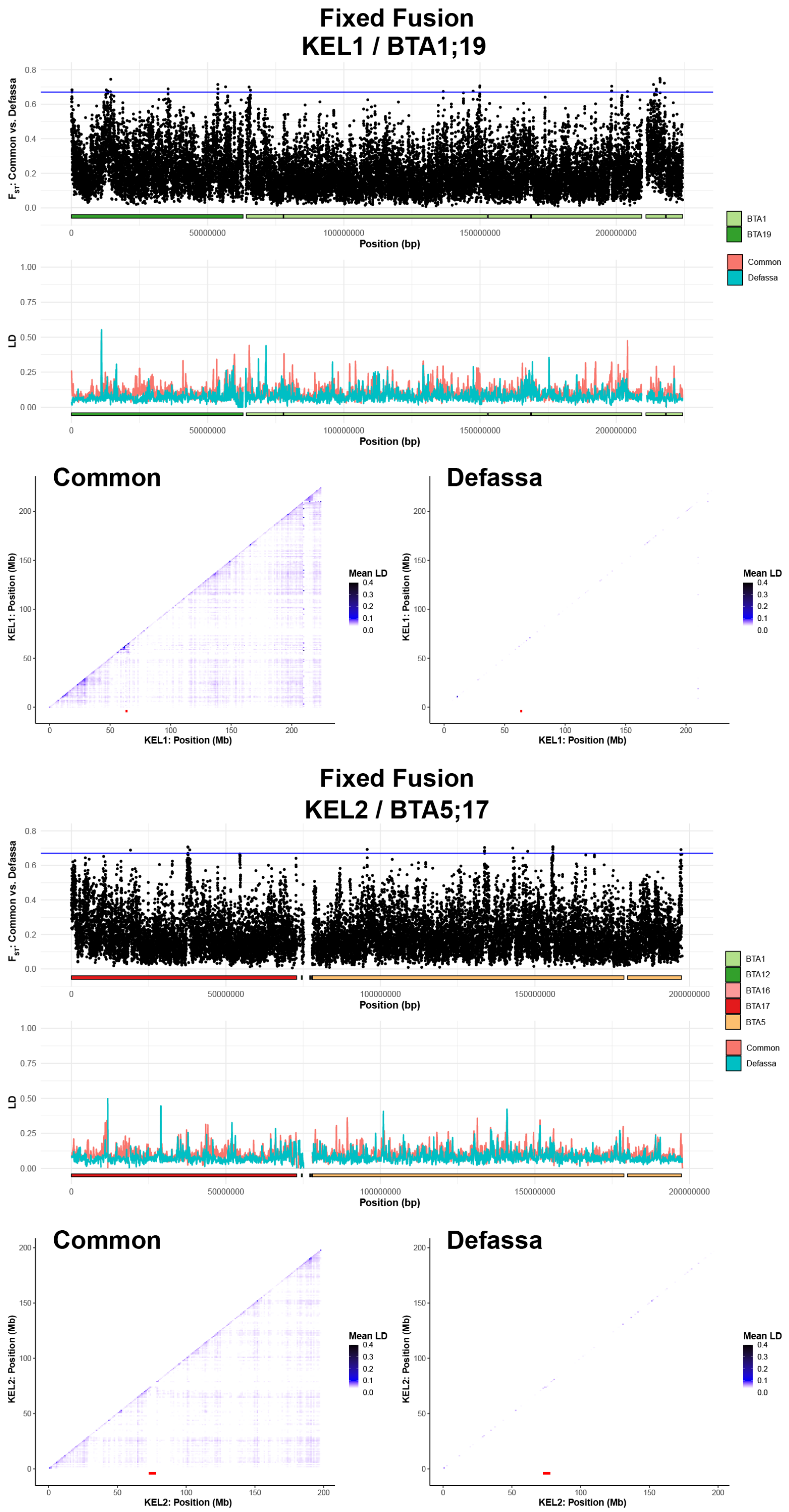


**Supplementary Fig. S12:** $F_{ST}$ between subspecies, mean LD in 100 Kb windows for each subspecies, and mean pairwise LD in 1 Mb windows for each subspecies across chromosomes KEL1 and KEL2 involved in fixed Rb fusions in waterbuck. Homology to cattle chromosomes (BTA) shown. Red line shows the putative centromeric region.

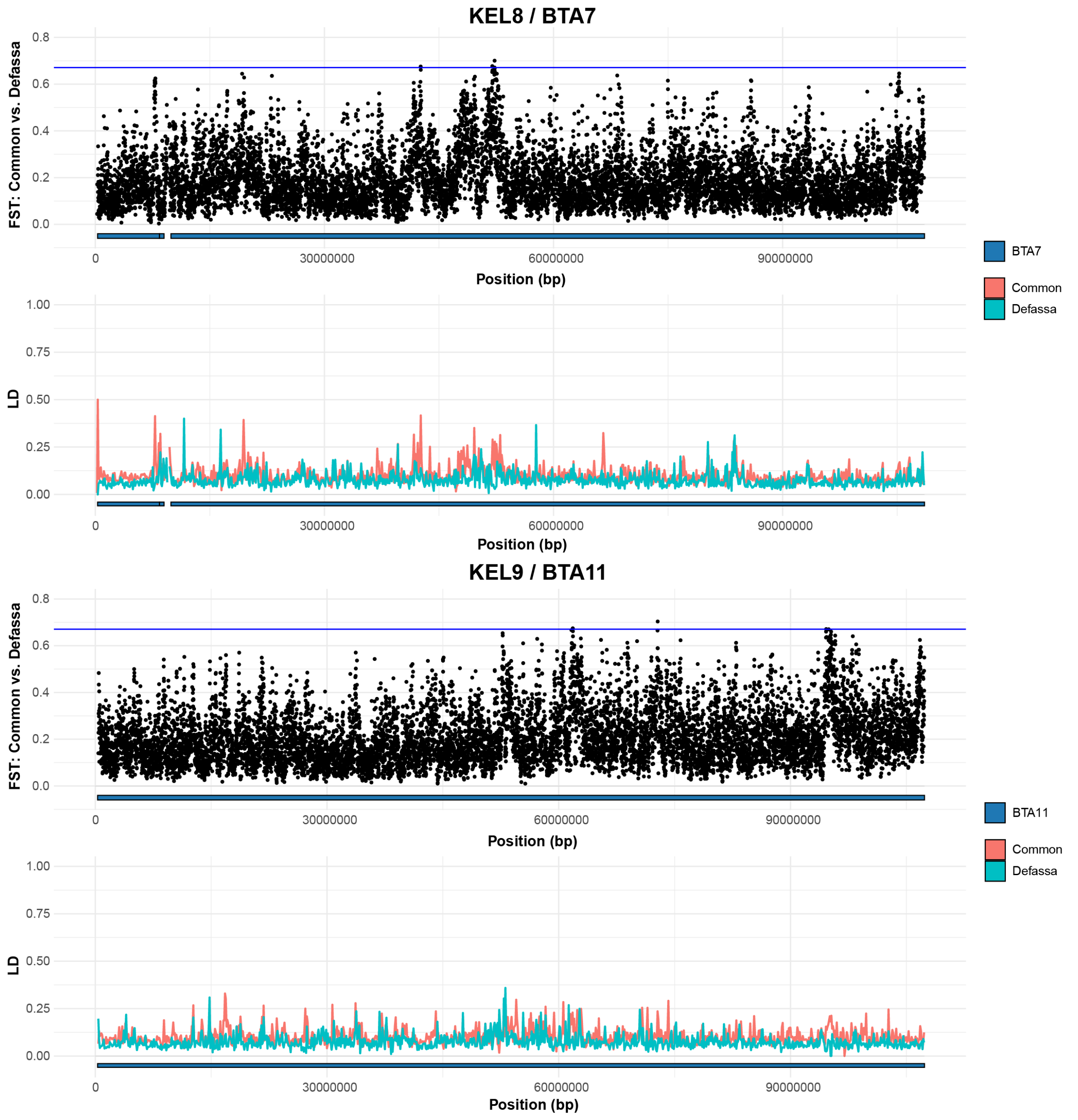


**Supplementary Fig. S13:** $F_{ST}$ between waterbuck subspecies and mean LD in 100 Kb windows for each subspecies across chromosomes KEL8 and KEL9 involved in the polymorphic Rb fusion. Homology to cattle chromosomes (BTA) shown.
